# *C. elegans* establishes germline versus soma by balancing inherited histone methylation

**DOI:** 10.1101/2020.01.22.914580

**Authors:** Brandon S. Carpenter, Teresa W. Lee, Caroline F. Plott, Jovan S. Brockett, Dexter A. Myrick, David J. Katz

**Author notes:** corresponding author: David J. Katz, Associate Professor, Department of Cell Biology, Room 443, Whitehead Biomedical Research Building, Emory University School of Medicine, Atlanta, GA 30322, USA.

## Abstract

Embryos undergo extensive reprogramming at fertilization to prevent the inappropriate inheritance of histone methylation. In *C. elegans,* this reprogramming is mediated by the H3K4me2 demethylase, SPR-5, and the H3K9 methyltransferase, MET-2. In contrast to this reprogramming, the H3K36 methyltransferase, MES-4, maintains H3K36me2/3 at germline genes between generations to help re-establish the germline. To determine whether the MES-4 germline inheritance system antagonizes *spr-5; met-2* reprogramming, we examined the interaction between these two systems. We find that the developmental delay of *spr-5; met-2* mutant progeny is associated with ectopic H3K36me2/3 and the ectopic expression of MES-4 targeted germline genes in somatic tissues. Furthermore, the developmental delay is dependent upon MES-4 and the H3K4 methyltransferase, SET-2. We propose that the MES-4 inheritance system prevents critical germline genes from being repressed by maternal *spr-5; met-2* reprogramming. Thus, the balance of inherited histone modifications is necessary to distinguish germline versus soma and prevent developmental delay.

## INTRODUCTION

In multicellular organisms, developmental cell fate decisions are established by tightly controlled spatial and temporal gene expression. This pattern of gene expression can be regulated by heritable modifications on the N-terminal tails of histone proteins (Jambhekar et al., 2019). For example, methylation of either lysine 4 or 36 on histone 3, (H3K4me and H3K36me) is generally associated with active transcription, whereas methylation of lysine 9 on the same histone (H3K9me) is commonly associated with transcriptional repression. These modifications are dynamically regulated by the specific and tightly controlled activity of histone modifying enzymes. For instance, mono- and di- methylation of lysine 4 on histone H3 (H3K4me1/2) are removed by the demethylase LSD1/KDM1A (Y. Shi et al., 2004; Y.-J. Shi et al., 2005). In the nematode *C. elegans*, populations of mutants lacking the LSD1 ortholog, SPR-5, become increasingly sterile over ~30 generations (Katz et al., 2009). Failure to erase H3K4me2 at fertilization between generations in *spr-5* mutants correlates with an accumulation of H3K4me2 and spermatogenesis gene expression across 30 generations, which leads to increasing sterility (Katz et al., 2009). These data demonstrate that H3K4me2 can function as an epigenetic transcriptional memory. More recently, we demonstrated that SPR-5 synergizes with the H3K9me2 methyltransferase, MET-2, to regulate maternal epigenetic reprogramming (Kerr et al., 2014). Progeny of mutants lacking both SPR-5 and MET-2 suffer from developmental delay and become completely sterile in a single generation (Kerr et al., 2014). These phenotypes are associated with synergistic increases in both H3K4me2 and candidate germline gene expression in somatic tissues (Kerr et al., 2014). Together this work supports a model in which SPR-5 and MET-2 are maternally deposited into the oocyte, where they reprogram histone methylation to prevent inherited defects.

Following fertilization, the *C. elegans* embryo separates germline versus somatic lineages progressively through a series of asymmetric divisions (Strome, 2005). To accomplish this, transcription factors coordinate with multiple histone modifications. For example, maternal deposition of PIE-1, a germline specific protein that asymmetrically segregates into germline blastomeres (P lineage cells), maintains the fate of germ cells by inhibiting POL-II elongation and preventing the ectopic expression of somatic genes (Batchelder et al., 1999; Mello et al., 1992; Seydoux et al., 1996). In the absence of transcription in the germline, the maternally provided H3K36me2/3 methyltransferase, MES-4, binds to a subset of germline genes that were previously expressed in the parental germline (Furuhashi et al., 2010; Rechtsteiner et al., 2010). These germline genes are recognized by MES-4 via H3K36me2/3 that was added in the parental germline by the transcription-coupled H3K36me2/3 methyltransferase, MET-1 (Kreher et al., 2018). MES-4 maintains H3K36me2/3 at these genes in the early embryo in a transcriptionally independent manner. Without maternally deposited MES-4, the germline cannot properly proliferate and animals are sterile (Capowski et al., 1991; Garvin et al., 1998). For the remainder of this study, we will refer to these MES-4 targeted genes as MES-4 germline genes, and the process through which the MES-4 germline inheritance system maintains these genes for re-activation in the offspring as bookmarking.

MES-4 bookmarking is antagonized in somatic tissues by transcriptional repressors and chromatin remodelers. For example, loss of the transcriptional repressors LIN-15B and LIN-35 at high temperatures leads to larval arrest (Petrella et al., 2011). This larval arrest can be suppressed by removing the MES-4 germline inheritance system (Petrella et al., 2011). Removing the MES-4 inheritance system also suppresses the somatic expression of germline genes in *lin-35* mutants (Wang et al., 2005). Similar to LIN-15B and LIN-35, loss of the chromatin remodelers MEP-1 and LET-418 causes somatic expression of germline genes and larval arrest (Unhavaithaya et al., 2002). This somatic expression of germline genes and larval arrest is also dependent upon the MES-4 germline inheritance system (Unhavaithaya et al., 2002). Together, these findings demonstrate that transcriptional repressors and the chromatin remodelers function in somatic tissues to antagonize H3K36 bookmarking by MES-4.

Recently, the repressive histone modification H3K9me2 has also been implicated in the somatic repression of germline genes (Rechtsteiner et al., 2019). Some germline genes have H3K9me2 enrichment at their promoters in somatic tissues (Rechtsteiner et al., 2019). Loss of LIN-15B reduces this enrichment of H3K9me2 leading to the ectopic accumulation of H3K36me3 at gene bodies in somatic tissues (Rechtsteiner et al., 2019). This raises the possibility that LIN-15B may antagonize the MES-4 germline inheritance in part through the repressive histone modification H3K9me2.

Despite the extensive knowledge of the systems that somatically antagonize the MES-4 germline inheritance system, it remains unclear why germline genes are bookmarked by H3K36 in the embryo. To address this major gap, we examined somatic development in progeny deficient in SPR-5 and MET-2 maternal reprogramming. Our previous work suggests that maternal *spr-5; met-2* reprogramming prevents the transgenerational inheritance of H3K4me2 by erasing this mark and coupling it to the acquisition of H3K9me2 between generations (Kerr et al., 2014). Here we show that H3K36me3 ectopically accumulates at MES-4 germline genes in the somatic tissues of *spr-5; met-2* double mutant progeny (hereafter referred to as *spr-5; met-2* progeny), and this accumulation correlates with the ectopic expression of these genes. In addition, we find that both the developmental delay and the ectopic expression of germline genes is rescued by knocking down MES-4 activity. These data provide comprehensive evidence that the ectopic expression of MES-4 targeted germline genes in somatic tissues causes developmental delay. In addition, we demonstrate that the severe developmental delay of *spr-5; met-2* progeny is rescued by the loss of the H3K4 methyltransferase SET-2. This finding suggests that the ectopic maintenance of the MES-4 germline inheritance system in *spr-5; met-2* progeny is driven by the inheritance of H3K4 methylation. Finally, by demonstrating that loss of maternal *spr-5; met-2* reprogramming unbalances the MES-4 germline inheritance system, our data suggest that H3K36 methylation bookmarking functions to antagonize *spr-5; met-2* maternal reprogramming. Thus, we propose that *C. elegans* balances three different histone modifications to distinguish between the competing fates of soma and germline.

## RESULTS

### Loss of *spr-5* and *met-2* causes a severe developmental delay

Previous observations from our lab indicated that progeny from *spr-5; met-2* mutants may develop abnormally (Kerr et al., 2014). To further characterize this phenotype, we synchronized embryos laid by N2, *spr-5*, *met-2*, and *spr-5; met-2* mutant hermaphrodites and monitored their development from hatching to adults. By 72 hours, all N2 progeny (469/469), most of the *spr-5* progeny (363/385), and many of *met-2* progeny (386/450) were fertile adults (Figure 1A, B, C, and E, Figure 1-figure supplement 1A-C). In contrast, *spr-5; met-2* progeny displayed a severe developmental delay, with none of the progeny (0/463) reaching adulthood by 72 hours (Figure 1D, E; and Figure 1-figure supplement 1D). The majority of *spr-5; met-2* progeny (371/463) resembled L2 larvae at 72 hours, while a smaller percentage of the population developed to later larval stages (42/463) (supplementary file 1). By seven days post synchronized lay, a small number of *spr-5; met-2* progeny (35/876) developed into adults and the majority (31/35) of these adults displayed protruding vulva (Pvl) (Figure 1-figure supplement 1E-G). All 35 of the *spr-5; met-2* mutant progeny that developed to adulthood were sterile.

**Figure 1.**
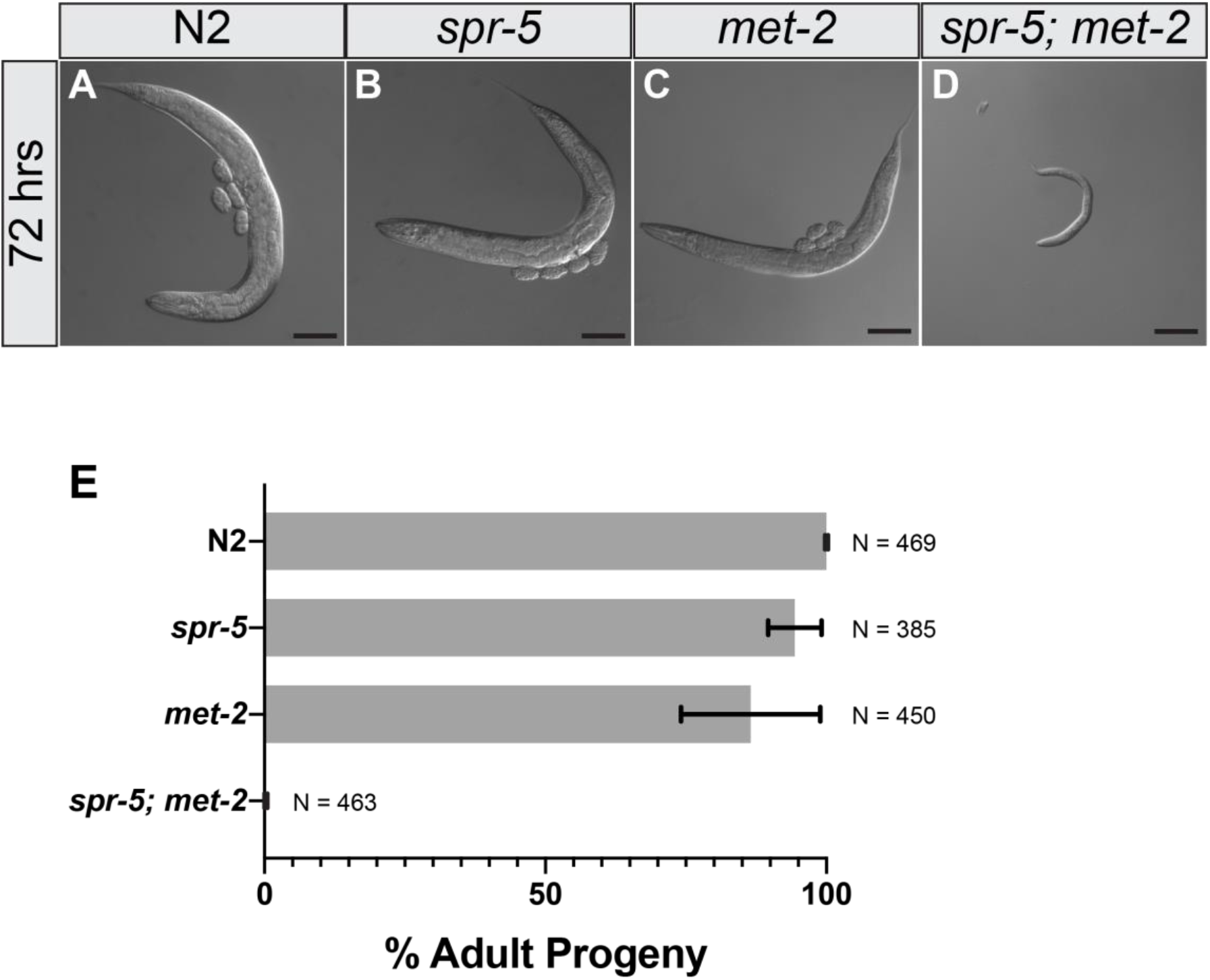
*spr-5; met-2* mutants display severe developmental delay. 10x Differential Interference Contrast (DIC) images of N2 (A), *spr-5* (B), *met-2* (C), and *spr-5; met-2* progeny (D) 72 hours post synchronized lay. Scale bar: 100μm. (E) Percentage of N2, *spr-5*, *met-2*, and *spr-5; met-2* progeny that reached the adult stage (% Adult Progeny) by 72 hours post synchronized lay. Error bars represent the standard deviation of the mean from three experiments. N= the total number of progeny 20-25 hermaphrodites scored over three experiments.

### MES-4 germline genes are ectopically expressed in *spr-5; met-2* mutant soma

Previously we showed that H3K4me2 is synergistically increased in *spr-5; met-2* progeny compared to *spr-5* and *met-2* single mutant progeny, and that this increase in H3K4me2 correlates with a synergistic increase in candidate germline gene expression in somatic tissues (Kerr et al., 2014). To test the extent to which germline genes are ectopically expressed in somatic tissues, we examined somatic expression genome-wide. To do this, we performed RNA-seq on *spr-5; met-2* L1 progeny compared to N2 (wild-type) L1 progeny. We chose to perform this analysis on L1 larvae because this stage immediately precedes the L2 larval delay that we observe in *spr-5; met-2* progeny (see Figure 1A, D). In addition, L1 larvae are composed of 550 somatic cells and two germ cells. Therefore, L1 larvae are primarily composed of somatic tissue. As a control, we also performed RNA-seq on L1 progeny from *spr-5* and *met-2* single mutants that were isolated from early generation animals, within the first five generations. These generations are well before the onset of sterility that we previously reported (Katz et al., 2009; Kerr et al., 2014).

We identified 778 differentially expressed transcripts in *spr-5; met-2* progeny compared to N2 (Figure 2-supplement figure 1A-B, Figure 2-supplement figure 2C, F, and supplementary file 2), many of which also overlap with genes differentially expressed in *spr-5* (159/413) and *met-2* single mutants (113/343) compared to N2 (Figure 2-supplement figure 1A-B, Figure 2-supplement figure 2A, B, D, E, and supplementary file 2). Gene Set Enrichment Analysis (GSEA) did not identify any categories of genes misexpressed in the *spr-5* or *met-2* single mutants. However, GSEA revealed that genes differentially expressed in *spr-5; met-2* progeny were significantly enriched (based on Combined Score, (Chen et al., 2013)) for biological processes and cellular components characteristic of the germline; including meiosis, P-granules and negative regulation of the cell cycle (Figure 2-supplement figure 1C-D). Many of the genes involved in these processes are expressed in the germline of the parental generation, bound by the H3K36 methyltransferase MES-4 in the early embryo, and marked by H3K36me2/3, independent of POL-II (Rechtsteiner et al., 2010). As a result, we were interested in the potential overlap with MES-4 targeted genes (referred to as MES-4 germline genes).

Rechtsteiner and colleagues identified approximately 200 MES-4 germline genes (Rechtsteiner et al., 2010). We reasoned that the absence of SPR-5 and MET-2 reprogramming may cause germline genes to be aberrantly targeted by MES-4 in the soma, leading to ectopic expression. To investigate this possibility, we examined the overlap between differentially expressed genes in *spr-5; met-2* L1 progeny and MES-4 germline genes. Out of 196 MES-4 germline genes, 34 overlapped with genes up-regulated in *spr-5; met-2* progeny compared to N2 (Figure 2A, hypergeometric test, p-value < 6.44e-20), while zero overlapped with genes down-regulated in *spr-5; met-2* progeny compared to N2 (Figure 2B). In addition, when we compared the log2 fold change (FC) in expression of all of the MES-4 germline genes in *spr-5*, *met-2*, and *spr-5; met-2* mutant progeny compared to N2, we observed that 108 of the MES-4 germline genes were synergistically increased in *spr-5; met-2* progeny compared to single mutant progeny (Figure 2C and supplementary file 3).

**Figure 2.**
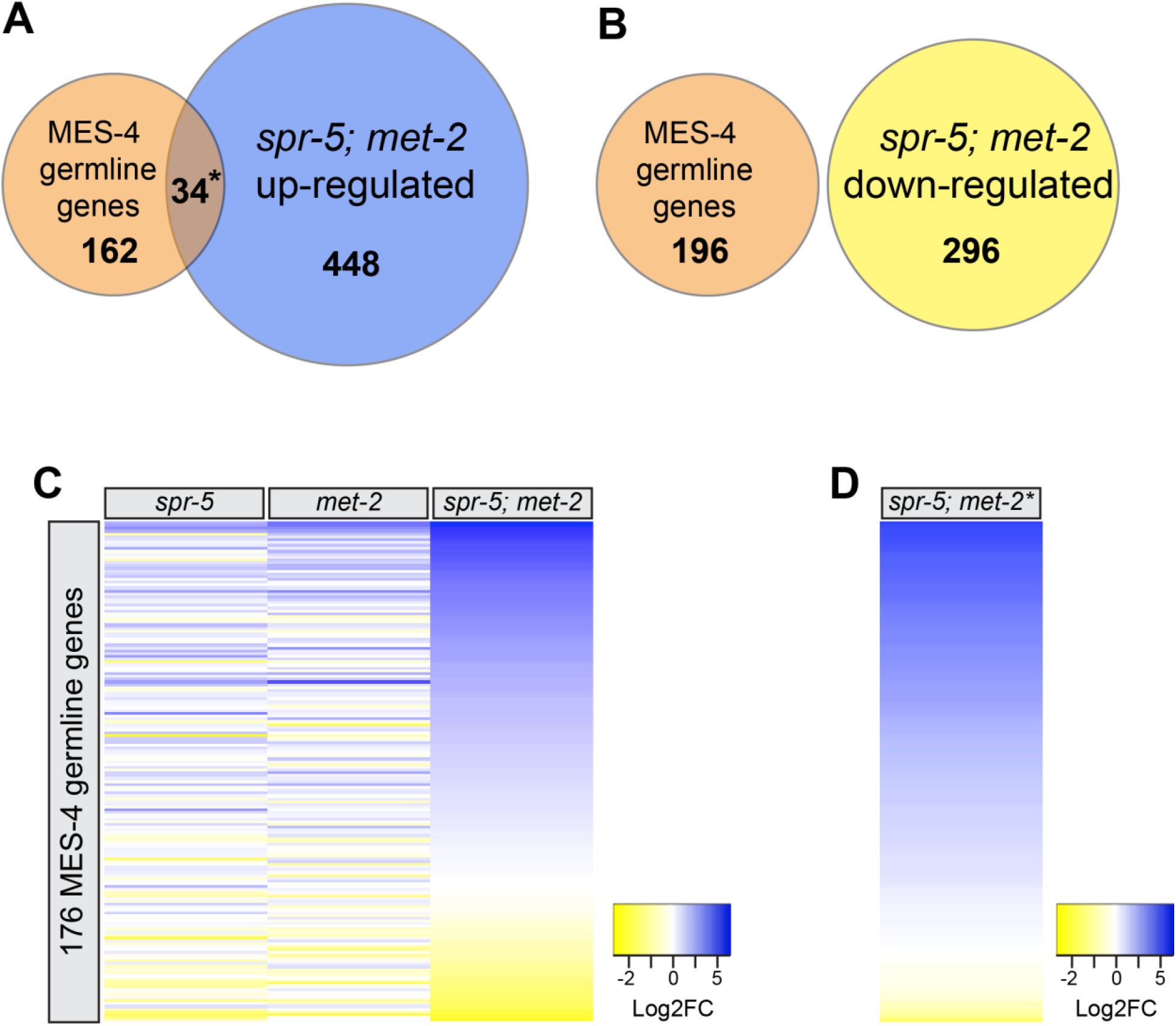
MES-4 germline genes are ectopically expressed in *spr-5; met-2* mutant soma. Overlap between MES-4 germline genes and genes up-regulated (A) and down-regulated (B) in *spr-5; met-2* L1 progeny. Significant over-enrichment in A was determined by the hypergeometric test (*p-value < 6.44e-20). (C) Heatmap of log2 fold change (FC) of 176 MES-4 germline genes in *spr-5, met-2* and *spr-5; met-2* mutants compared to N2. (D) Heatmap of log2FC of 176 MES-4 germline genes for the repeat *spr-5; met-2* RNA-seq experiment, with three additional biological replicates of *spr-5; met-2\** mutants compared to N2 (see methods). Log2FC values are represented in a yellow to blue gradient and range from −2 to 5. Yellow represents genes with negative log2FC values and blue represents genes with positive log2FC values compared to N2. The remaining 21 MES-4 germline genes were not included because they do not have an expression value in one or more of the data sets (*spr-5*, *met-2,* or *spr-5; met-2*).

During this initial RNA-seq analysis we had to genotype every *spr-5; met-2* L1 because the balancer chromosome that was available did not completely balance *spr-5*. As a result, the RNA-seq was performed using a low-input sequencing technique (see methods). However, during the course of the experiments, a new balancer became available that completely balances *spr-5*. This enabled us to repeat the *spr-5; met-2* RNA-seq experiments using standard amounts of RNA. In the repeat *spr-5; met-2* RNA-seq experiment, we identified significantly more differentially expressed genes compared to N2 (4223 vs. 778 in the initial low-input analysis). However, despite the larger number of differentially expressed genes, MES-4 germline genes remained similarly enriched. Out of 196 MES-4 germline genes, 112 overlapped with genes up-regulated in *spr-5; met-2* progeny compared to N2 (Figure 2-supplemental figure 3A, hypergeometric test, P-value < 1.20e-54, and supplementary file 2), while only two overlapped with genes down-regulated in *spr-5; met-2* progeny compared to N2 (Figure 2-supplemental figure 3B). We also compared the log2FC in expression of all of the MES-4 germline genes in *spr-5; met-2* mutant progeny compared to N2. This analysis revealed that the MES-4 germline genes were similarly overexpressed in *spr-5; met-2* progeny compared to N2 (Figure 2D and supplementary file 3). Interestingly, while MES-4 germline genes were enriched in both *spr-5; met-*2 RNA-seq experiments, there were some differences in the specific MES-4 germline genes that were overexpressed, and the extent to which they were overexpressed (Figure 2-supplemental figure 3C and supplementary file 3).

### smFISH confirmation of MES-4 germline gene expression in *spr-5; met-2* mutant soma

To confirm that MES-4 germline genes are somatically expressed in *spr-5; met-2* L1 progeny, we performed single molecule fluorescent *in situ* hybridization (smFISH) on two MES-4 germline targets, *htp-1* and *cpb-1* (Figure 3). Both of these genes were amongst the genes that were ectopically expressed in *spr-5; met-2* L1 progeny compared to N2 L1 progeny. In N2 L1 larvae, *htp-1* (Figure 3A-C, insets) and *cpb-1* (Figure 3G-I, insets) were restricted to the two primordial germ cells, Z2 and Z3, which go on to form the entire adult germline. This confirms that these transcripts are confined to the germline as expected. In contrast, in *spr-5; met-2* progeny *htp-1* was ectopically expressed throughout the soma (Figure 3D-F). This expression pattern is similar to what we observed with the ubiquitously expressed subunit of RNA polymerase II, *ama-1* (Figure 3-supplement figure 1A-L), which was unchanged in our RNA-seq analysis. *cpb-1* wa*s* also ectopically expressed in *spr-5; met-2* progeny, though the ectopic expression was not as ubiquitous as *htp-1* (Figure 3J, L).

**Figure 3.**
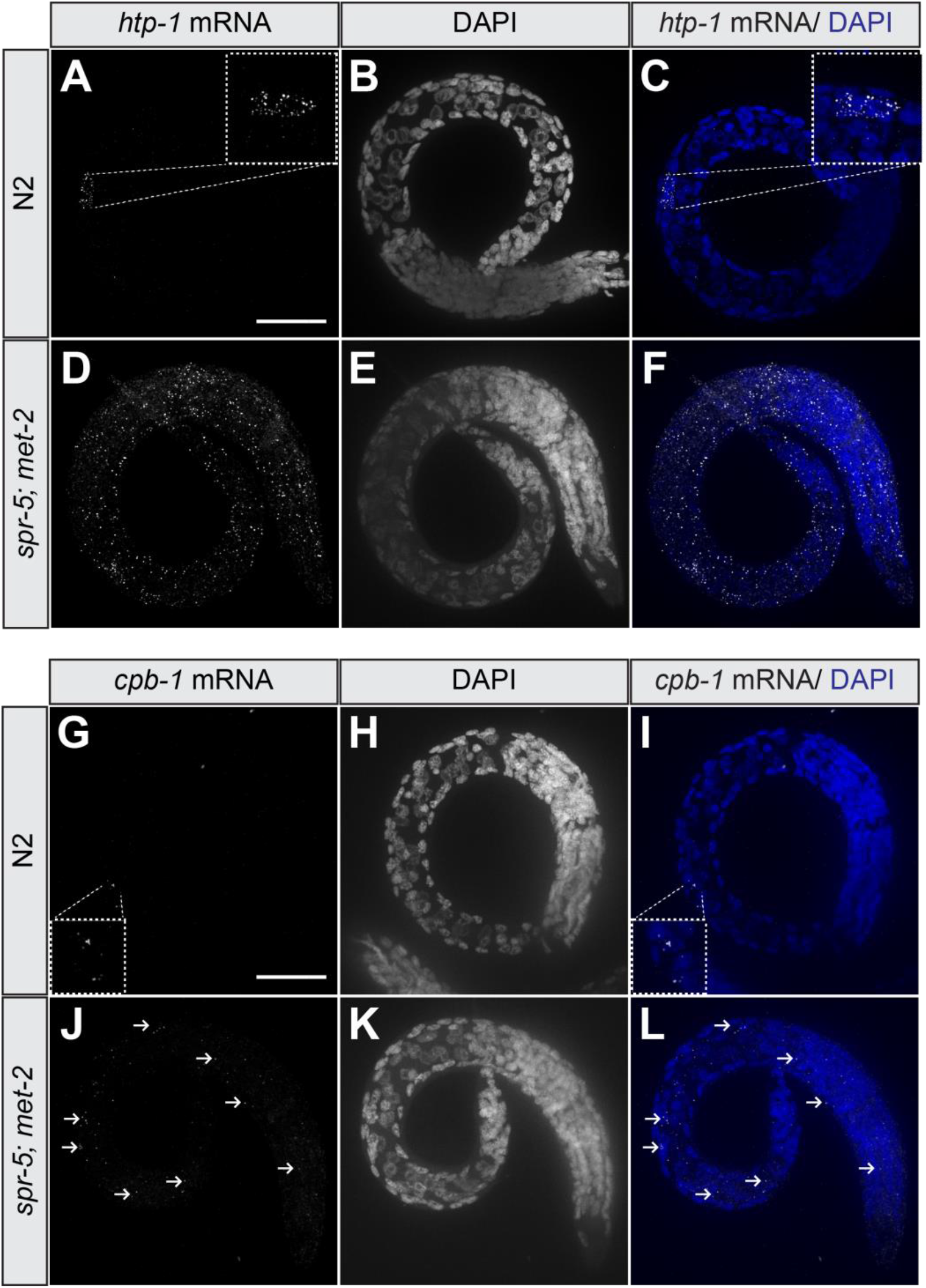
*spr-5; met-2* L1 progeny ectopically express MES-4 germline genes in multiple somatic tissues. 40x smFISH images of *htp-1* (A, C, D, F) and *cpb-1* (G, I, J, L) endogenous mRNAs in N2 (A-C, G-I) and *spr-5; met-2* (D-F, J-L) L1 progeny. DAPI was used as a nuclear marker (B, C, E, F, H, I, K, L). Insets are high magnification images of the germ cells, Z2 and Z3, in N2 L1 progeny. Arrows (J, L) denote ectopic *cpb-1* mRNA foci in somatic cells. Scale bar 40μm.

### MES-4 germline genes maintain ectopic H3K36me3 in *spr-5; met-2* mutants

To test whether MES-4 germline genes that are ectopically expressed in the soma of *spr-5; met-2* progeny also ectopically maintain H3K36me3, we performed H3K36me3 ChIP-seq. MES-4 germline genes have low levels of H3K36me3 in N2 L1 progeny. However, compared to N2 L1 progeny, *spr-5; met-2* L1 progeny displayed increased enrichment for H3K36me3 across gene bodies at MES-4 germline genes (Figure 4A, B; Figure 4-supplement figure 1A, B, Q-R). For example, the MES-4 germline genes *cpb-1*, *T05B9.1, Y18D10A.11*, and *fbxa-101,* that are ectopically expressed in our RNA-seq analysis, showed increased levels of H3K36me3 in *spr-5; met-2* progeny (Figure 4J-M; Figure 4-supplement figure 1J-M) compared to N2 progeny (Figure 4C-F; Figure 4-supplement figure 1C-F). As a control, we examined H3K36me3 enrichment at genes that are not affected in *spr-5; met-2* progeny. These control genes include: *ceh-13*, a gene enriched in hypodermal and ventral nerve chord in L1 progeny (Figure 4G, N; Figure 4-supplement figure 1G, N), *ama-1*, a subunit of RNA polymerase II that is expressed ubiquitously (Figure 4H, O; Figure 4-supplement figure 1H, O), and *act-1*, a ubiquitously expressed actin related protein (Figure 4I, P; Figure 4-supplement figure 1I, P). Each of these control genes displayed similar H3K36me3 enrichment in both *spr-5; met-2* and N2 L1 progeny (compare Figure 4N-P and Figure 4-supplement figure 1N-P to Figure 4G-I and Figure 4-supplement figure 1G-I, respectively).

**Figure 4.**
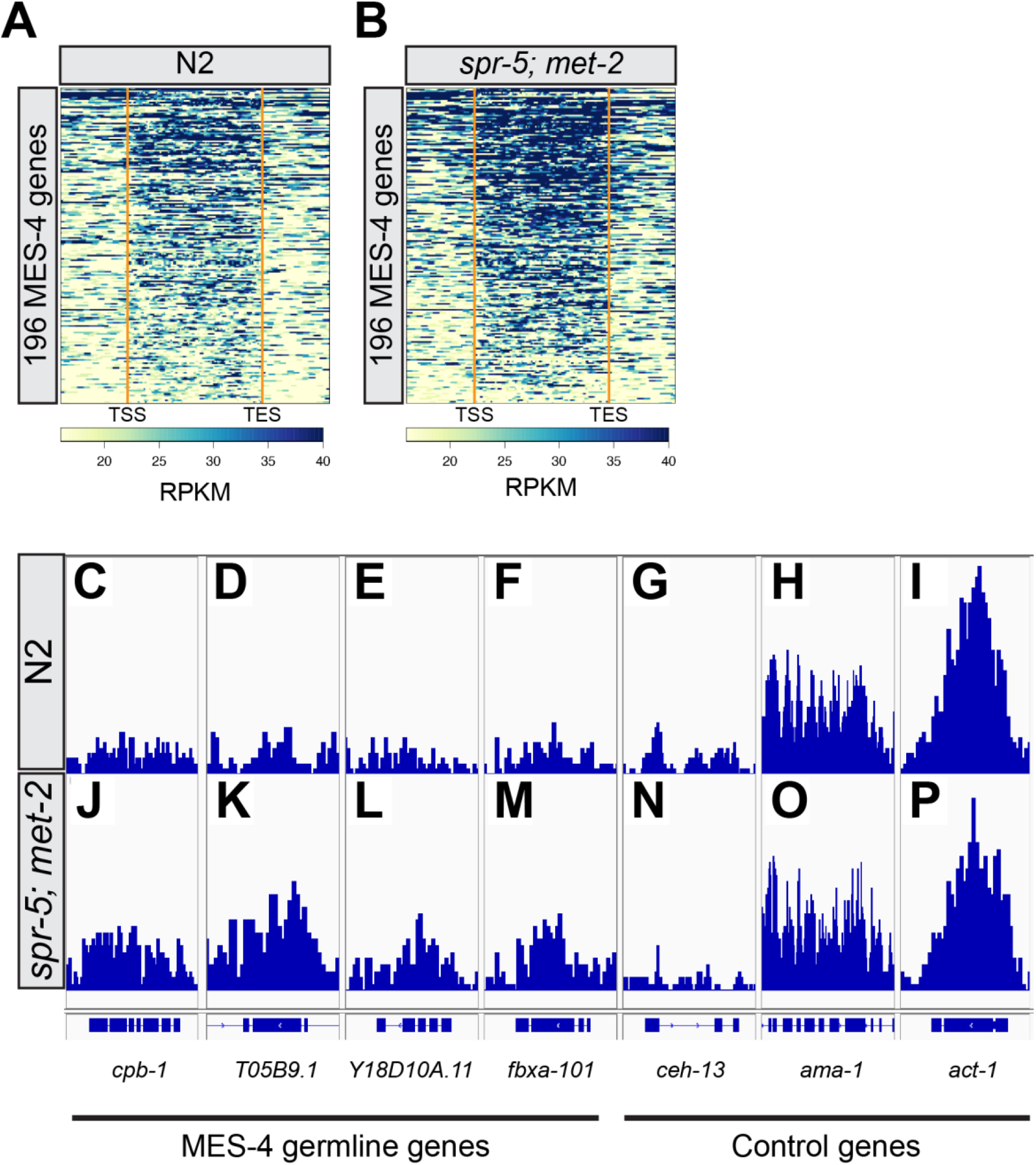
MES-4 germline genes display ectopic H3K36me3 in *spr-5; met-2* L1 progeny. Heatmap of H3K36me3 ChIP-seq reads normalized to reads per kilobase million (RPKM) over the gene bodies of 196 MES-4 germline genes in wild-type (N2) (A) versus *spr-5; met-2* (B) L1 progeny (second replicate in Figure S5). Gene bodies were pseudoscaled to 1kb with 500bp borders separated by orange bars that represent the transcriptional start site (TSS) and transcriptional end site (TES). Integrative Genome Viewer (IGV) image of H3K36me3 ChIP-seq reads normalized to RPKM at MES-4 germline genes (C-F, J-M) and control genes (G-I, N-P) in N2 (C-I) versus *spr-5; met-2* (J-P) L1 progeny. RPKM IGV windows were scaled between 0 and 202 RPKM for all genes.

### MES-4 germline genes display H3K9me2 at their promoter peaks

Recent work discovered that some germline specific genes contain H3K9me2 peaks at their promoters in N2 L1 progeny (Rechtsteiner et al., 2019). This finding implicates H3K9me2 enrichment at promoters of germline genes as being a critical component for repressing germline genes in somatic tissues. If SPR-5 and MET-2 are functioning to prevent MES-4 germline genes from being ectopically expressed in somatic tissues, we would expect MES-4 germline genes that are ectopically expressed in the somatic tissues of *spr-5; met-2* progeny to normally continue to be targeted by H3K9 methylation in these tissues. To examine this possibility, we re-analyzed L1 stage H3K9me2 Chip-seq data from Rechsteiner et al. 2019 (Rechtsteiner et al., 2019). This re-analysis showed that many of the MES-4 germline genes were enriched for H3K9me2 at their promoters (Figure 4-supplement figure 2A), including the majority of MES-4 germline genes that were ectopically expressed in the soma of *spr-5; met-2* progeny (Figure 4-supplement figure 2B). For example, the MES-4 germline genes *cpb-1*, *T05B9.1*, *Y18D10A.11*, and *fbxa-101*, that were misexpressed somatically and accumulated ectopic H3K36me3 in the somatic tissues of *spr-5; met-2* progeny, had H3K9me2 peaks at their promoters (Figure 4-supplement figure 2C-F). In contrast, our control genes *ceh-13*, *ama-1*, and *act-1*, that were not misexpressed, were also not enriched for H3K9me2 at their promoters (Figure 4-supplement figure 2G-I).

### Knocking down MES-4 rescues ectopic expression of germline genes in *spr-5; met-2* mutant soma

To test whether the ectopic expression of MES-4 germline genes in *spr-5; met-2* progeny is dependent on the ectopic H3K36me3, we examined whether the expression of these genes was dependent upon MES-4. We performed quantitative real-time PCR (qRT-PCR) on L1 progeny from *spr-5; met-2* hermaphrodites fed control (L4440) RNAi versus *mes-4* RNAi (Figure 5). For this analysis, we selected candidate MES-4 germline genes that were ectopically expressed and displayed an ectopic H3K36me3 peak in *spr-5; met-2* L1 progeny compared to N2 L1 progeny. Consistent with our RNA-seq analysis, all nine of the candidate MES-4 germline genes that we examined were ectopically expressed >2 fold in *spr-5; met-2* L1 progeny compared to control (Figure 5). Strikingly, the ectopic expression of the nine MES-4 candidate germline genes was dependent upon MES-4. Nine out of nine of these genes were significantly decreased in L1 progeny from *spr-5; met-2* hermaphrodites treated with *mes-4* RNAi (Figure 5; unpaired t-test, p-value < 0.001), and all but one (*T05B9.1*) were reduced to levels that were similar to N2 L1 progeny.

**Figure 5.**
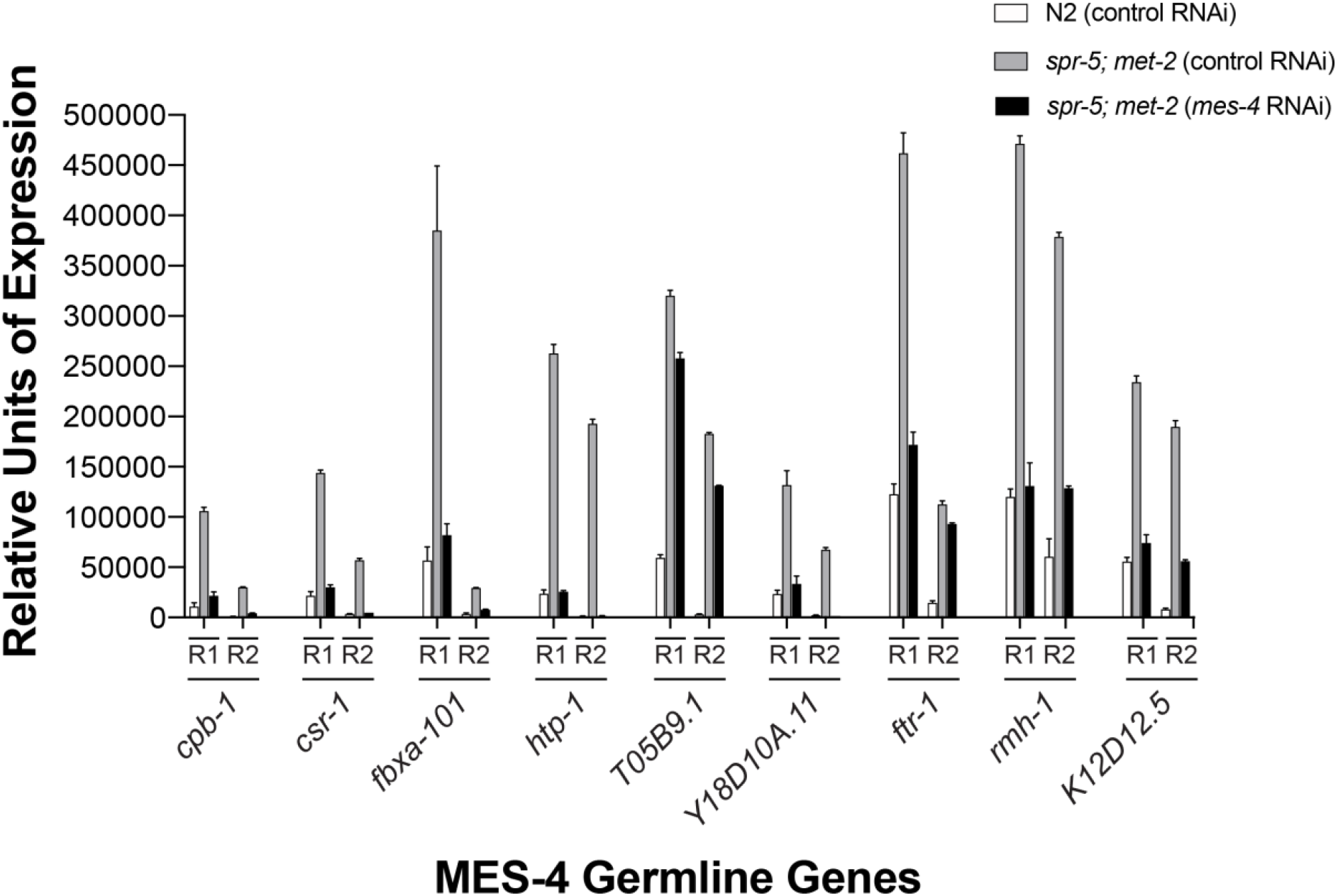
Knocking down MES-4 rescues ectopic expression of MES-4 germline genes in *spr-5; met-2* L1 progeny. Quantitative RT-PCR showing the relative units of expression for nine MES-4 germline genes (*cpb-1, csr-1*, *fbxa-101*, *htp-*1, *T05B9.1*, *Y18D10A.11*, *ftr-1*, *rmh-1*, *K12D12.5*) in L1 progeny of *spr-5; met-2* hermaphrodites fed either control L4440 RNAi (grey bars) or *mes-4* RNAi (black bars) versus N2 fed control L4440 RNAi (white bars). Relative units of expression from two biological replicates (R1 and R2) were calculated for each gene by averaging triplicate RT_PCR reactions and normalizing to a control gene, *ama-1*. Error bars represent the standard error of the mean (SEM) for triplicate RT_PCR reactions. For all nine genes, *mes-4* RNAi significantly reduced the relative expression of *spr-5; met-2* compared to *spr-5; met-2* fed L4440 control RNAi (unpaired t-test, p-value <0.001).

### Knocking down MES-4 rescues developmental delay in *spr-5; met-2* progeny

To test whether the developmental delay phenotype that we observe in *spr-5; met-2* progeny is also dependent on the ectopic expression of MES-4 germline genes, we fed *spr-5; met-2* hermaphrodites *mes-4* RNAi and monitored their progeny for 72 hours after a synchronized lay. If the developmental delay is dependent upon the ectopic expression of MES-4 germline genes, it should be suppressed when this ectopic expression is eliminated via *mes-4* RNAi. By 72 hours, all of the N2 control progeny from hermaphrodites fed either L4440 control (1089/1089) or *mes-4* (1102/1102) RNAi were adults (Figure 6A, B, I). Also, consistent with our previous observations, all but one of the *spr-5; met-2* mutant progeny (729/730) from hermaphrodites fed control RNAi remained in the L2-L3 larval stages (Figure 6E, I). In contrast, most of *spr-5; met-2* progeny (569/618) from hermaphrodites fed *mes-4* RNAi developed to adults (Figure 6F, I, unpaired t-test, p<0.0001). Though as expected, these animals remained sterile due to the *mes-4* RNAi preventing any germline formation.

**Figure 6.**
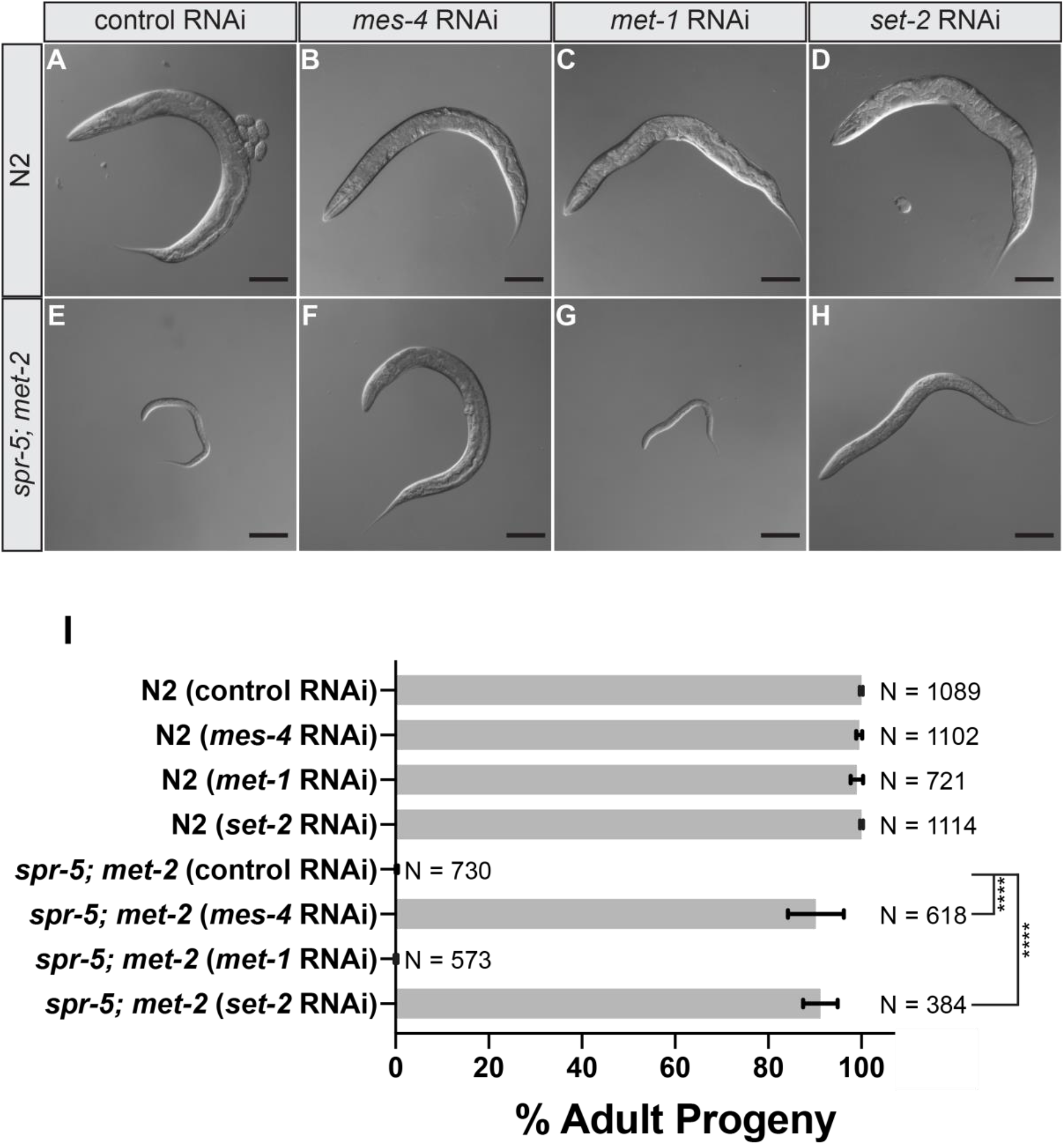
Knocking down MES-4 rescues developmental delay in *spr-5; met-2* progeny. DIC images of N2 (A-D) or *spr-5; met-2* (E-H) progeny from hermaphrodite parents treated with control (L4440 vector only) RNAi (A, E), *mes-4* RNAi (B, F), *met-1*RNAi (C, G), or *set-2* RNAi (D, H) 72 hours post synchronized lay. Scale bar: 100μm. (I) Quantification of the number of progeny (represented as % Adult Progeny) from A-H that made it to adults by 72 hours. Error bars represent the standard deviation of the mean from two or three experiments. N represents the total number of progeny from 30-40 hermaphrodites scored across independent experiments. (unpaired t-test, **** represent a p-value <0.0001).

MET-1 mediates transcription-coupled H3K36me3 in the adult germline and in later stage transcriptionally active embryos (Furuhashi et al., 2010; Kreher et al., 2018; Rechtsteiner et al., 2010). If the ectopic somatic expression of MES-4 germline genes is caused by the transcription independent maintenance of H3K36me3 via MES-4, we would expect the developmental delay to be independent of MET-1. To determine whether the developmental delay phenotype in *spr-5; met-2* progeny is dependent on MET-1, we monitored the development of progeny from *spr-5; met-2* hermaphrodites fed *met-1* RNAi for 72 hours after a synchronized lay. In contrast to the suppression that we observed upon *mes-4* RNAi, *met-1* RNAi had no effect. By 72 hours, none of the *spr-5; met-2* mutant progeny (0/573) from hermaphrodites fed *met-1* RNAi developed to adults, and these animals were indistinguishable from *spr-5; met-2* progeny fed control RNAi (Figure 6G, I). Additionally, *met-1* RNAi itself had no effect on developmental delay, as nearly all of the N2 progeny (715/721) from hermaphrodites fed *met-1* RNAi developed to adults (Figure 6C, I).

### Knocking down SET-2 rescues *spr-5; met-2* developmental delay

If the developmental delay of *spr-5; met-2* mutants is caused by ectopic inheritance of H3K4me2 driving the expression MES-4 germline genes in somatic tissues, we would expect the developmental delay of *spr-5; met-2* progeny to be dependent upon the H3K4 methyltransferase, SET-2. To test this, we monitored the development of progeny of *spr-5; met-2* hermaphrodites fed *set-2* RNAi for 72 hours after a synchronized lay. Identical to N2 progeny from hermaphrodites fed control RNAi, *set-2* RNAi had no effect on the development of wild-type animals, as all of the N2 progeny from hermaphrodites fed *set-2* RNAi developed to adults by 72 hours (1114/1114) (Figure 6D, I). However, in contrast to *spr-5; met-2* progeny fed control RNAi that were developmentally delayed, most of the progeny from *spr-5; met-2* hermaphrodites fed *set-2* RNAi developed to adults (347/384) (Figure 6H, I, unpaired t-test, p<0.0001).

### *spr-5; met-2* progeny acquire transgene silencing in somatic tissues

The somatic expression of MES-4 germline genes involved in germline transgene silencing (Figure S2C-D) raises the possibility that the somatic tissues in *spr-5; met-2* progeny may acquire the ability to silence transgenes, a function normally restricted to germline cells. To test this, we examined the somatic expression of an extrachromosomal multicopy *let-858* transgene that is normally silenced in the germline by both transcriptional and posttranscriptional germline silencing mechanisms (Kelly and Fire, 1998). This analysis was performed in *spr-5; met-2* mutant L2 larvae that were undergoing developmental delay. In N2, most of the L2 progeny (117/132) expressed ubiquitous high levels of LET-858::GFP throughout the entire soma (Figure 7A, B, and K), while the remaining progeny (15/132) expressed what we describe as a “faint” level of expression (Figure 7C, D, and K). In contrast, almost none of the *spr-5; met-2* L2 progeny (2/87) displayed high level of transgene expression that is comparable to the high level seen in most N2 progeny (Figure 7E, F, and K). Instead, the majority of *spr-5; met-2* progeny (64/87) had faint LET-858::GFP expression that is comparable to the faint expression observed in N2 progeny (Figure 7G, H, and K). The remaining 21 *spr-5; met-2* progeny had no LET-858::GFP expression. Because less than 50% of progeny inherited the *let-858* transgene, we normalized the percentage of progeny scored as “off” for LET-858::GFP to the presence of the *let-858* transgene based on genotyping for *gfp* (see methods). For N2, all of the 60 L2 progeny that were scored as “off” failed to inherit the *let-858* transgene, indicating that the transgene is never silenced in N2 progeny (data not shown). However, for *spr-5; met-2* L2 progeny, eight out of 50 progeny that were scored as “off” inherited the *let-858* transgene indicating that the *let-858* transgene can be completely silenced in some *spr-5; met-2* progeny. After normalization for the transgene inheritance, we observed that 21/87 of *spr-5; met-2* progeny displayed no visible expression of the LET-858::GFP transgene (Figure 7I, J, and K).

**Figure 7.**
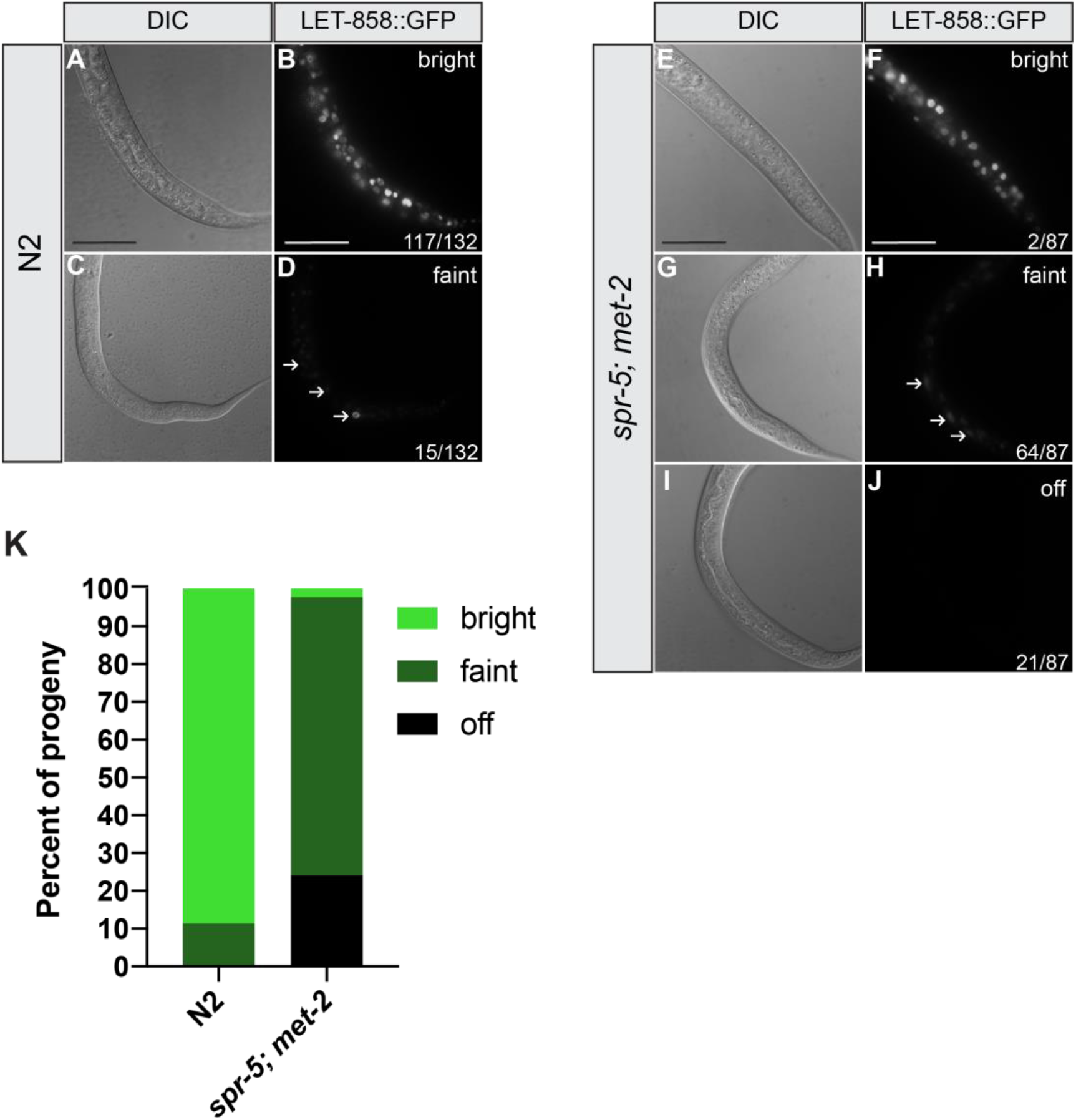
*let-858* transgene silencing in the soma of *spr-5; met-2* mutants. 40x differential contrasting interference (DIC) (A, C, E, G, and I) and immunofluorescent (B, D, F, H, and J) images of N2 (A-D) and *spr-5; met-2* (E-J) L2 progeny. Arrows denote faint expression of LET-858::GFP. Scale bar: 50 μm. Scale bar is the same for all panels. Percentage of animals where the expression level of LET-858::GFP was scored as either bright expressing (bright green, representative shown in panel B and F), faint expressing (dark green, representative shown in panel D and H), or not expressing (black, representative shown in panel J) in N2 (N=132) versus *spr-5; met-2* (N=87) progeny (K). The quantification represents the percentages of LET-858::GFP expressing progeny from two independent experiments. To control for the segregation of the *let-858* transgene, progeny scored as “off” were normalized for the presence of the *let-858* transgene in animals as detected by PCR (see methods).

## DISCUSSION

### *spr-5; met-2* maternal reprogramming prevents developmental delay by restricting ectopic MES-4 bookmarking

SPR-5 and MET-2 act maternally to reprogram histone methylation and prevent the transcriptional state of the parent from being inappropriately transmitted to the offspring (Kerr et al., 2014). In this study, we find that the loss of SPR-5 and MET-2 maternal reprogramming led to a severe developmental delay that was associated with the ectopic expression of MES-4 germline genes in somatic tissues. This finding raises the possibility that SPR-5 and MET-2 reprogramming blocks the ectopic expression of MES-4 germline genes by preventing the accumulation of MES-4 dependent H3K36me3 at a subset of germline genes in somatic tissues. Consistent with this possibility, most of the MES-4 germline genes were increased in *spr-5; met-2* mutants compared to N2 in L1 larvae. Using smFISH, we also confirmed the somatic expression of two ectopically expressed MES-4 germline genes, *htp-1* and *cpb-1*. While *htp-1* mRNA was ectopically detected in many somatic tissues, the ectopic expression of *cpb-1* mRNA was more restricted, suggesting that the extent of ectopic expression is dependent upon the locus.

In the absence of SPR-5 and MET-2 reprogramming, MES-4 germline genes accumulated ectopic H3K36me3 in the soma. This suggests that without SPR-5 and MET-2 reprogramming, MES-4 ectopically maintains H3K36me3 at these genes in somatic tissues. Of note, we observe a low level of H3K36me3 at germline genes in the somatic tissues of N2 progeny. It is unclear why there is a low level of H3K36me3 normally in somatic tissues in N2 animals. Nevertheless, the absence of transcription associated with this low level of H3K36me3 indicates that an increased level of H3K36me3 is necessary to cause ectopic transcription.

If the ectopic maintenance of H3K36me3 in the soma of *spr-5; met-2* mutant progeny is causing the developmental delay, then removal of MES-4 should rescue the ectopic expression and developmental delay. Indeed, we find that the removal of MES-4 rescued both the ectopic transcription of MES-4 germline genes in the soma of *spr-5; met-2* progeny, and the developmental delay. Importantly, removing the transcriptionally coupled H3K36me3 methyltransferase, MET-1, did not rescue the developmental delay. Thus, the developmental delay that we observe in *spr-5; met-2* progeny is only dependent on transcriptionally independent H3K36me3. Taken together, our data provide comprehensive evidence that the developmental delay of *spr-5; met-2* progeny is caused by the ectopic expression of MES-4 germline genes.

### How does an ectopic transcriptional program interfere with developmental timing?

How does the ectopic expression of germline genes interfere with somatic tissues to cause developmental delay? One possibility is that the ectopic expression of MES-4 germline genes causes the soma to take on germline character. To begin to address this, we asked whether *spr-5; met-2* double mutants can silence an extrachromosomal array in somatic tissues. The silencing of extrachromosomal arrays is normally restricted to the germline (Kelly and Fire, 1998). However, we find that *spr-5; met-2* progeny acquired some ability to silence an extrachromosomal multicopy array in somatic cells. Consistent with this finding, loss of the somatic repressor LIN-35 also results in the somatic silencing of a GFP transgene (Wang et al., 2005). Intriguingly, in the *spr-5; met-2* mutant RNA-seq we detected the ectopic expression of RNA-dependent RNA polymerase genes (e.g. *rrf-1* and *gld-2*) as well as genes involved in the RNAi effector complex (e.g. *hrde-1* and *ppw-1*). These pathways have previously been implicated in gene silencing (Buckley et al., 2012; Sijen et al., 2001; Tijsterman et al., 2002). Thus, similar to what has been found in *lin-35* mutants (Wang et al., 2005), it is possible that the somatic silencing of the transgene in *spr-5; met-2* mutants is due to the induction of the germline small RNA pathway. In a reciprocal fashion, heritable silencing via small RNAs requires MET-2. This provides a further indication that the chromatin and small RNA pathways may be functioning together (Lev et al., 2017).

Normally in L1 larvae, the primordial germ cells, Z2 and Z3, are arrested at the G2/M checkpoint (Fukuyama et al., 2006). In the *spr-5; met-2* mutant RNA-seq, we also detected the ectopic expression of genes involved in the negative regulation of proliferation and the cell cycle, as well as G2/M checkpoint genes. Thus, it is also possible that ectopic expression of germline genes normally expressed only in Z2 and Z3 contributes to the developmental delay through the ectopic activation of germline cell cycle control. Regardless, the silencing of the extrachromosomal multicopy array suggests that the somatic tissues in *spr-5; met-2* progeny make functional proteins that can perform some germline functions. We propose that either ectopic germline transcription, or an ectopic germline function resulting from this ectopic transcription, interferes with the ability of somatic cells to properly enact their transcriptional program. This background noise delays the proper adoption of cell fate, leading to an overall delay in the development of the tissue.

### A model for how the inheritance of histone methylation is balanced to specify germline versus soma

By linking maternal *spr-5; met-2* reprogramming to the MES-4 germline inheritance system, our data provide a rationale for the existence of MES-4 bookmarking, through the following model. In the germline, transcriptional elongation is blocked by PIE-1, which segregates to germline blastomeres during embryogenesis (Batchelder et al., 1999; Mello et al., 1992; Seydoux et al., 1996) (Figure 8A, B). In the soma of the early embryo, there is also very little transcription, because the bulk of zygotic transcription does not begin until approximately the 60-cell stage. This stage is just prior to when the primordial germ cells, Z2 and Z3, are specified (Sulston et al., 1983). Thus, in *C. elegans,* germline versus soma is largely specified without transcription.

**Figure 8:**
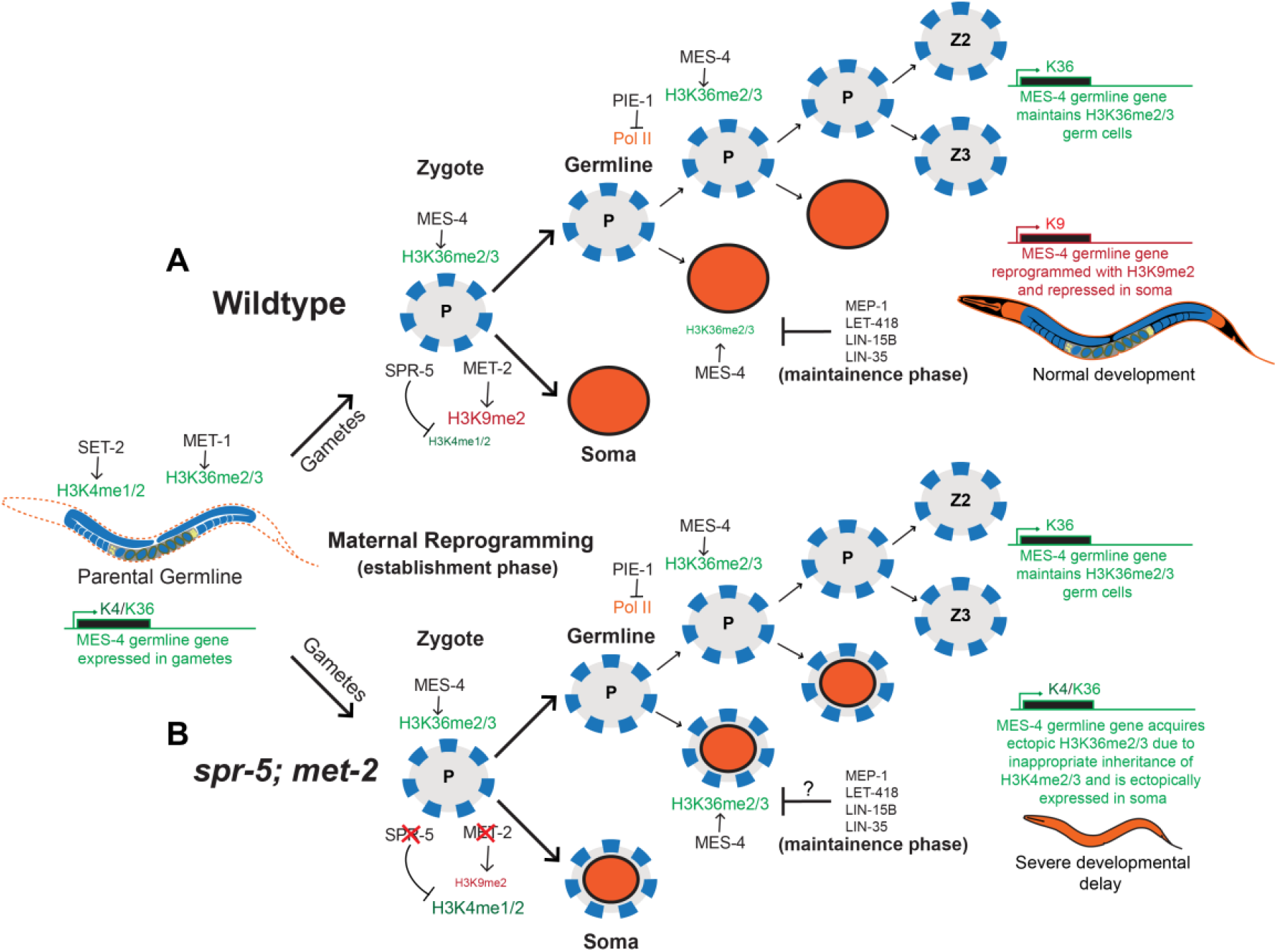
A model for how maternal reprogramming of inherited histone methylation helps to specify germline versus soma. During development, SET-2 and MET-1 add transcriptionally coupled H3K4me1/2 and H3K36me2/3 to germline expressed genes in the parental germline, respectively. (A) At fertilization, these germline expressed genes undergo maternal epigenetic reprogramming (establishment phase) by SPR-5 and MET-2 to remove H3K4me1/2 and add H3K9me1/2. In the germline blastomeres of the embryo, PIE-1 prevents global transcription by inhibiting POL-II. In the absence of transcription, MES-4 maintains H3K36me2/3 at MES-4 germline genes that have acquired transcriptionally coupled H3K36me2/3 in the previous germline. This enables these genes to avoid being repressed by maternal *spr-5; met-2* reprogramming and ensures that these genes remain bookmarked for re-expression once the germline begins to proliferate later in development. In addition, multiple systems, such as LIN-15B and LIN-35, as well as MEP-1 and LET-418, function in somatic tissues to further antagonize H3K36 bookmarking by MES-4 (maintenance phase). (B) Without SPR-5 and MET-2 maternal reprogramming, H3K4me1/2 is inappropriately inherited in somatic tissues, allowing MES-4 to ectopically add H3K36me2/3 at these germline genes. This leads to ectopic expression of MES-4 germline genes in somatic tissues and a severe developmental delay. Orange circles represent somatic cells, Grey circles outlined in blue dashed-lines represent germ cells, and orange circles outlined in dashed-blue lines depict somatic cells that ectopically express MES-4 germline genes. P-lineage, germline blastomeres, are indicated by the letter P, and the primordial germline cells are indicated by Z2 and Z3.

During normal maternal reprogramming, SPR-5 and MET-2 are deposited into the oocyte. At fertilization they facilitate the reprogramming of previously expressed genes from an active chromatin state to a repressed chromatin state by removing H3K4me2 and adding H3K9me2 (Kerr et al., 2014) (Figure 8A). This reprogramming is necessary to prevent the transcriptional memory of the previous generation from being inappropriately propagated to the progeny. Genes epigenetically reprogrammed by SPR-5 and MET-2 include ubiquitously expressed genes and germline expressed genes, a subset of which are MES-4 germline genes. The MES-4 germline genes are subsequently targeted by the transcription independent H3K36 methyltransferase, MES-4, to maintain H3K36me3 in the germ lineage during embryogenesis (Figure 8A, B). We propose that H3K36 methylation bookmarking antagonizes the repression caused by the erasure of H3K4me2 and the addition of H3K9me2. Without the transcription independent maintenance of inherited H3K36 methylation from the mother to antagonize this repression, the germline fails to proliferate and animals are sterile (Capowski et al., 1991; Garvin et al., 1998). The failure to proliferate and sterility caused by loss of MES-4 is presumably because germline genes that are targeted by MES-4 fail to reactivate, though it has yet to be demonstrated. Thus, our data suggest that the MES-4 bookmarking system is necessary for critical germline genes to bypass the global epigenetic reprogramming that occurs at fertilization to prevent transgenerational inheritance. We refer to this initial phase of filtering inherited histone methylation at fertilization as the establishment phase.

Following this establishment phase, a maintenance phase is required to propagate this initial pattern of histone methylation throughout embryogenesis. The MES-4 bookmarking system is localized primarily to the primordial germ cells in later embryonic development (Furuhashi et al., 2010; Rechtsteiner et al., 2010). This concentration of MES-4 to the germline helps to maintain MES-4 bookmarking for germline specification later in embryonic development. However, MES-4 is also present in somatic tissues (Furuhashi et al., 2010; Rechtsteiner et al., 2010). This makes the system vulnerable to the ectopic maintenance of MES-4 bookmarking in the soma. For this reason, multiple systems, such as LIN-15B and LIN-35, as well as MEP-1 and LET-418, function in somatic tissues to restrict H3K36 bookmarking by MES-4 to the germline. Thus, in the maintenance phase, the balance of MES-4 and the systems that somatically antagonize MES-4, maintain the histone methylation pattern that is initiated during the establishment phase. Taken together, we propose that SPR-5, MET-2 and MES-4 carefully balance the inheritance of three different histone modifications, H3K4, H3K9, and H3K36 methylation, to ensure the proper specification of germline versus soma in the absence of transcription.

The model that we have proposed makes the following two predictions. First, MES-4 germline genes should normally be targeted for continued silencing by H3K9me2 in somatic tissues. It has recently been shown that a subset of germline specific genes contain H3K9me2 at their promoters in somatic tissues (Rechtsteiner et al., 2019). We re-examined the H3K9me2 ChIP-seq dataset from this work and found that the MES-4 germline genes that are ectopically expressed in the somatic tissues of *spr-5; met-2* progeny also displayed unique H3K9me2 promoter peaks. This confirms that MES-4 germline genes are normally repressed by H3K9me2 in somatic tissues.

The second prediction from our model is that the ectopic inheritance of H3K4 methylation at MES-4 germline genes overwhelms the somatic repression systems. Despite the presence of these transcriptional repressor complexes and chromatin remodelers to antagonize MES-4 bookmarking in the soma, loss of *spr-5; met-2* maternal reprogramming results in the somatic expression of MES-4 germline genes. This suggests that the failure to add H3K9me2, as well as the inappropriate retention of H3K4me2, results in a chromatin environment that is permissive for the ectopic maintenance of H3K36me3 in the soma, even in the presence of the systems that repress MES-4 bookmarking somatically (Figure 8B). If this is the case, then the developmental delay in *spr-5; met-2* progeny should also be dependent upon the activity of the H3K4 methyltransferase. We find that knocking down SET-2, the H3K4me1/2 methyltransferase, rescued the developmental delay that we observe in *spr-5; met-2* progeny. These findings suggest that the inheritance of ectopic H3K4 methylation enables the ectopic accumulation of MES-4 dependent H3K36me3, and the subsequent ectopic expression of MES-4 germline genes in somatic tissues of *spr-5; met-2* progeny. It is not entirely clear how the subsequent ectopic maintenance of H3K36me3 facilitates ectopic expression. However, it should be noted that there were some differences in which MES-4 germline genes were ectopically expressed between our two *spr-5; met-2* RNA-seq experiments. This stochasticity is consistent with H3K36me3 being permissive, rather than instructive, for transcription. If this is the case, it is doubtful that the *spr-5; met-2* developmental delay is caused by the inappropriate expression of any single MES-4 germline gene. Rather, it is likely that the developmental delay is caused by either the inappropriate expression of multiple MES-4 germline genes, or the ectopic activation of the MES-4 germline program.

### Conservation of maternal epigenetic reprogramming between invertebrates and vertebrates

Epigenetic reprogramming at fertilization is a problem that all sexually reproducing organisms must solve (Lee and Katz, 2020). Thus, it is possible that the mechanisms of epigenetic reprogramming may be conserved. Along with the Heard lab, we previously demonstrated that progeny from mice that maternally lack the vertebrate SPR-5 homolog, KDM1A/LSD1, ectopically maintain the expression of germline genes in the embryo. This causes embryonic lethality at the two-cell stage (Ancelin et al., 2016; Wasson et al., 2016). Similarly, maternal loss of the MET-2 ortholog SETDB1 or the MES-4 ortholog NSD1 in mice results in early embryonic lethality (J. Kim et al., 2016; Rayasam et al., 2003). Together these results underscore the developmental importance of properly regulating histone methylation between generations and raise the possibility that the mechanism we have uncovered is conserved in mammals.

The model that we have proposed may also help explain the mechanism underlying patients harboring mutations in various histone-modifying enzymes. Recent genome sequencing has revealed that several neurodevelopmental disorders are caused by mutations in histone modifying enzymes (extensively reviewed by (J.-H. Kim et al., 2017). These include mutations in: 1) the H3K36 methyltransferase *Setd2* and the H3K27 demethylase *Kdm6a,* which cause Kabuki Syndrome (Lederer et al., 2012), 2) the human ortholog of *spr-5, Lsd1,* which causes a Kabuki-like Syndrome (Chong et al., 2016; Tunovic et al., 2014), and 3) the H3K36 methyltransferase *Nsd1* which causes Sotos Syndrome (Kurotaki et al., 2002). Similar to what we observed in *spr-5; met-2* mutant progeny, many of the human patients with mutations in these histone modifying enzymes suffer from global developmental delay. Based on our model, it is possible that the developmental delay in these patients may be caused by the failure to properly regulate histone methylation during critical developmental transitions. The resulting inappropriate inheritance of histone methylation could result in the ectopic expression of a developmental program in an inappropriate tissue, leading to background noise and developmental delay.

## MATERIALS AND METHODS

### Strains

All *Caenorhabditis elegans strains* were grown and maintained at 20° C under standard conditions, as previously described (Brenner, 1974). The *C. elegans spr-5 (by101)(I)* strain was provided by R. Baumeister. The N2 Bristol wild-type (WT) strain was provided by the *Caenorhabditis* Genetics Center. The *met-2 (n4256)(III) strain* was provided by R. Horvitz. The *hT2 [bli-4(e937)let-?(q782)qls48] (I;III)* balancer strain was used to maintain *spr-5 (by101)(I); met-2 (n4256)(III)* double-mutant animals as heterozygotes. Because the *hT2 [bli-4(e937)let-?(q782)qls48](I;III)* balancer allele does not extend completely to the *spr-5* locus on chromosome I, the F0 animals used to generate F1 *spr-5; met-2* progeny were cloned out and genotyped to confirm the presence of the *spr-5 (by101)(I)* allele. For genotyping, single animals were picked into 5-10ul of lysis buffer (50mM KCl, 10mM Tris-HCl (pH 8.3), 2.5mM MgCl2, 0.45% NP-40, 0.45% Tween-20, 0.01% gelatin) and incubated at 65°C for 1 hour followed by 95°C for 30 minutes. PCR reactions were performed with AmpliTaq Gold (Invitrogen) according to the manufacturer’s protocol and reactions were resolved on agarose gels (see supplementary file 5 for primer sequences). Before completing this study we acquired the FX30208 *tmC27 [unc-75(tmls1239*)](I) from the *Caenorhabditis* Genetics Center that completely covers the *spr-5* locus on chromosome I. The *qC1 [qls26 (lag2::GFP + rol-6(su1006)](III)* strain was obtained from W. Kelly and crossed to *met-2 (n4256)(III)* to maintain *met-2(n4256)(III)* as heterozygotes. The *spr-5 (by101)(I)*/*tmC27[unc-75(tmls1239)](I); met-2 (n4256) (III*)/qC1 [qls26 (lag2::gfp+ rol-6(su1006))](III) strain was then re-created for this study to maintain *spr-5 (by101)(I); met-2 (n4256)(III)* double-mutant animals as balanced heterozygotes. The LET-858::GFP (*pha-1*(e2123ts)(III); *let-858::gfp* (ccEx7271)) (Kelly and Fire, 1998) transgenic strain used in somatic transgene silencing assays was acquired from W. Kelly.

### Scoring developmental delay

*C. elegans* adult hermaphrodites were allowed to lay embryos for 2-4 hours and then removed in order to synchronize the development of progeny. Progeny were then imaged and scored for development to the adult stage at either 72 hours or seven days after synchronized lay, depending on the experiment.

### RNA sequencing and analysis

Total RNA was isolated using TRIzol reagent (Invitrogen) from 200-250 starved L1 larvae born at room temperature (21°C - 22°C) overnight in M9 Buffer. Due to difficulty in isolating large numbers of *spr-5; met-2* double-mutant progeny from the *hT2 [bli-4(e937)let-?(q782)qls48](I;III)* balancer strain, we submitted total RNA to the Genomic Services Laboratory (GSL) (HudsonAlpha, Huntsville, Alabama) for low input RNA-seq services. This service utilizes the Ovation RNA-Seq System V2 kit (Nugen) for initial RNA amplification prior to library preparation and sequencing (Illumina HiSeq v4, 50bp paired-end reads). For each genotype, 2 biological replicates were obtained. During the course of these experiments, the FX30208 *tmC27 [unc-75(tmls1239*)](I) balancer became available from the *Caenorhabditis* Genetics Center. This balancer completely covers the *spr-5* locus on chromosome I. Using this balanced strain, we performed a repeat *spr-5; met-2* RNA-seq experiment with three additional biological replicates of *spr-5; met-2* versus N2 L1 progeny. We submitted the total RNA from new replicates of the repeat RNA-seq to Georgia Genomics and Bioinformatics Core (University of Georgia, Athens, Georgia) for standard Poly-A RNA-seq services (Illumina Nextseq, 50bp paired-end reads). Downstream quality control and analysis were performed identically for both RNA-seq experiments. For both the low-input and repeat standard RNA-seq, sequencing reads were checked for quality using FastQC (Wingett and Andrews, 2018), filtered using Trimmomatic (Bolger et al., 2014), and remapped to the *C. elegans* transcriptome (ce10, WS220) using HISAT2 (D. Kim et al., 2015). Read count by gene was obtained by FeatureCounts (Liao et al., 2014). Differentially expressed transcripts for the low-input RNA-seq experiment (significance threshold, Wald test, p-value < 0.05) and the repeat RNA-seq experiment (significance threshold, Wald test, p-adj < 0.05) were determined using DESEQ2 (v.2.11.40.2) (Love et al., 2014). Transcripts per million (TPM) values were calculated from raw data obtained from FeatureCounts output. Subsequent downstream analysis was performed using R with normalized counts and p-values from DESEQ2 (v.2.11.40.2). Heatmaps were produced using the ComplexHeatmap R Package (Gu et al., 2016). Data was scaled and hierarchical clustering was performed using the complete linkage algorithm. In the linkage algorithm, distance was measured by calculating pairwise distance. Volcano plots were produced using the EnhancedVolcano package (v.0.99.16). Additionally, Gene Ontology (GO) Pathway analysis was performed using the online platform WormEnrichr (Chen et al., 2013; Kuleshov et al., 2016). Rechtsteiner and colleagues identified 214 MES-4 germline genes (Rechtsteiner et al., 2010). 17 of these genes are pseudogenes that we were unable to convert from Ensembl transcript IDs to RefSeq mRNA accession, and another gene was duplicated, so we removed those genes. This leaves 196 MES-4 germline genes that we used for our analysis. An additional heatmap comparison of differentially expressed genes between *spr-5*, *met-2*, and *spr-5; met-2* progeny compared to N2 progeny was generated in Microsoft Excel using log2 fold change values from the DESEQ2 analysis. Because transcript isoforms were ignored, we discuss the data in terms of “genes expressed” rather than “transcripts expressed”.

### Chromatin immunoprecipitation sequencing and analysis

Chromatin immunoprecipitation (ChIP) experiments were performed as described by Katz and colleagues (Katz et al., 2009). Briefly, 600 starved L1 larvae born at room temperature (21°C - 22°C) overnight in M9 Buffer were collected, frozen in liquid nitrogen, and stored at −80°C prior to homogenization. Frozen pellets were disrupted by a glass Dounce homogenizer, fixed with formaldehyde (1% final concentration), and quenched with glycine. ChIP samples were processed with a Chromatin Immunoprecipitation Assay Kit (Millipore), according to manufacturer’s instructions. Samples were sonicated using a Diagenode Bioruptor UCD-200 at 4°C on the “high” setting for a total of 30min with a cycle of 45sec on and 15sec off. A total of 12.5μL (5ug) H3K36me3 antibody (cat. 61021; Active motif) was used for immunoprecipitation. The Genomic Services Laboratory (GSL) (HudsonAlpha, Huntsville, Alabama) performed library preparation and sequencing (Illumina HiSeq v4, 50bp single-end reads). Reads were checked for quality using FastQC (Wingett and Andrews, 2018) and remapped to the *C. elegans* transcriptome (ce10, WS220) using Bowtie2 (Langmead and Salzberg, 2012; Langmead et al., 2009) under default parameters. bamCoverage in deepTools2 (Ramírez et al., 2016) was used to generate bigwig coverage tracks in 50bp bins, with blacklisted regions from McMurchy et al. 2017 excluded, using the following parameters: -bs 50, --normalizeUsing RPKM (McMurchy et al., 2017). MACS2 (Feng et al., 2012; Zhang et al., 2008) default parameters were used to call peaks and create bedgraph files for sequenced and mapped H3K36me3 ChIP samples and input DNA samples with the following adjustments to account for H3K36me3 broader domains: Broad-cutoff =0.001. Blacklisted regions from McMurchy et al. 2017 were excluded for this analysis (McMurchy et al., 2017). Using published H3K36me3 modMine Chip-chip called broad peaks (modENCODE_3555) from N2 L1 larvae as a guide, we then merged called broad peaks within 1200bp using Bedtools: MergeBED (Quinlan and Hall, 2010). H3K9me2 bedgraph files used in our analysis were from a published dataset (Rechtsteiner et al., 2019). Integrative Genome Viewer (IGV) was used to visualize H3K36me3 reads normalized to reads per kilobase millions (RPKM) and H3K9me2 reads normalized to 15 million reads (genome wide coverage of H3K9me2; (Rechtsteiner et al., 2019).

### RNAi methods

RNAi by feeding was carried out using clones from the Ahringer library (Kamath and Ahringer, 2003). Feeding experiments were performed on RNAi plates (NGM plates containing 100 ug/ml ampicillin, 0.4mM IPTG, and 12.5ug/ml tetracycline). F0 worms were placed on RNAi plates as L2 larvae and then moved to fresh RNAi plates 48hrs later where they were allowed to lay embryos for 2-4 hrs. F0 worms were then removed from plates and sacrificed or placed in M9 buffer overnight so that starved L1 progeny could be isolated for quantitative PCR (qPCR). F1 progeny were scored 72hrs after the synchronized lay for developmental progression. For each RNAi experiment, *pos-1* RNAi was used as a positive control. Each RNAi experiment reported here *pos-1* RNAi resulted in >95% embryonic lethality, indicating that RNAi plates were optimal.

### Real-time expression analysis

Total RNA was isolated using TRIzol reagent (Invitrogen) from synchronized L1s born at room temperature (21°C – 22°C). cDNA synthesis and qPCR were carried out as described (Kerr et al. 2014). A total of two biological replicates were performed and for both biological replicates experiments were performed in triplicate and normalized to *ama-1* mRNA expression (see supplementary file 5 for primer sequences).

### Differential interference contrast microscopy

Worms were immobilized in 0.1% levamisole and placed on a 2% agarose pad for imaging at either 10x or 40x magnification. 40x DIC images were overlaid together using Adobe photoshop to generate high resolution images of whole worms.

### Single Molecule Fluorescent *in situ* Hybridization (smFISH)

Quasar 570 labeled smFISH probe sets for *htp-1* and *cpb-1* were designed using Stellaris Probe Designer (Biosearch) (see supplementary file 4 for probe sequences). The *htp-1* smFISH probes were designed using the complete 1,059nt *htp-1* protein-coding sequence. Likewise, The *cpb-1* smFISH probes were designed using the complete 1,683nt *cpb-1* protein-coding sequence. In addition, an smFISH fluorescent probe set for *ama-1* was purchased from the DesignReady catalog (cat#: VSMF-6002-5, Biosearch). Synchronized L1 larvae for smFISH were obtained by bleaching 300-500 gravid hermaphrodites and allowing embryos to hatch overnight on 6cm NGM plates lightly seeded OP50 bacteria. L1 larvae were then washed into 1.5ul Eppendorf tubes using nuclease-free M9 buffer. Fixation and hybridization steps followed the Stellaris RNA FISH protocol for *C. elegans* adapted from RAJ lab protocol (Raj and Tyagi, 2010). In brief, we resuspended L1 larvae in fixation buffer (3.7% formaldehyde in 1 x PBS) for 15 minutes at room temperature then transferred tubes to liquid nitrogen. Samples were thawed in water and placed on ice for 20 minutes. In our hands, we obtain better fluorescent signal by freeze cracking L1 larvae. Following fixation, L1 Larvae were resuspended in 70% EtOH and stored at 4°C for 24-48 hours. For all probe sets, we incubated L1 larvae in 100ul hybridization buffer (containing 10% formamide and 125nm probe) for 4 hours at 37°C. After hybridization, samples were washed in wash buffer at 37°C for 30 minutes, incubated in 50ng/mL DAPI in wash buffer at 30°C for 30 minutes, washed once in 2x SSC for 2 minutes at room temperature, and mounted in Vectashield mounting medium. Mounted slides were imaged immediately using a 100x objective on a spinning-disk confocal Nikon-TiE imaging system. Images were captured using NIS-Elements software (Nikon) and ImageJ (NIH, http://imagej.nih.gov/ij/) was used for viewing. ImageJ maximum projection was used to project z-stack images to a single plane. The fluorescent intensity of smFISH dots were > 2-fold above background as expected (Ji and van Oudenaarden, 2012).

### LET-858::GFP transgene silencing assay

First generation *spr-5; met-2* hermaphrodites were crossed to *let-858* transgenic males to generate *spr-5/+*; *met-2/+*; *let-858::gfp* animals. The let-858 transgene is an extrachromosomal multicopy *let-858* array (Kelly and Fire, 1998). From these animals we generated *spr-5; met-2; let-858::gfp* animals and scored them for somatic expression of LET-858::GFP using a standard stereoscope. L2 progeny from N2 and *spr-5; met-2* progeny expressing the LET-858::GFP transgene were scored as “bright” (High level ubiquitous expression), faint (barely visible and ubiquitous expression), or off (no expression). Because less than 50% of progeny inherit the *let-858* transgene, we normalized the percentage of progeny scored as “off” for LET-858::GFP to presence of the *let-858* transgene based on genotyping for *gfp.* For N2, 0 out of 60 progeny that were scored as “off” failed to inherit the *let-858* transgene indicating that the transgene is never silenced in N2 progeny (data not shown, see supplementary file 5 for primer sequences). For *spr-5; met-2* progeny, 8 out of 50 progeny that were scored as “off” inherited the *let-858* transgene indicating that the *let-858* transgene is completely silenced in some *spr-5; met-2* progeny (data not shown).

## ACKNOWLEDGEMENTS

We are grateful to members of the Katz lab, as well as T. Caspary and C. Bean, for their helpful discussion and critical reading of the manuscript; M. J. Rowley for advice on bioinformatics; S. Strome and L. Petrella for helpful discussion and experimental advice; D. Lerit, R. Deal, V. Corces, and W. Kelly for the generous sharing of reagents and equipment. We thank R. Horvitz, W. Kelly, and the *Caenorhabditis* Genetics Center (funded by NIH P40 OD010440) for strains.

## Author Contributions

B.S.C. and D.J.K. conceived and designed the study and wrote the manuscript. B.S.C. performed experiments under the direction of D.J.K. B.S.C and D.J.K. analyzed data and interpreted results, with help from T.W.L. J.S.B. initially performed *set-2* RNAi rescue experiment and C.F.P. initially performed the *let-858* transgene silencing experiment. D.A.M. helped with RNA-seq analysis. All authors discussed the results.

## Data availability

Raw and processed genomic data has been deposited with the Gene Expression Omnibus (www.ncbi.nlm.nih.gov/geo) under accession code GSE143839.

## Supplementary Material

Figure Supplements:

Figure 1-figure supplement 1

Figure 2-figure supplements 1, 2, and 3

Figure 3-figure supplement 1

Figure 4-figure supplements 1 and 2

Supplementary files 1-6

Source Data files 1-3

---

**Figure 1 - supplement figure 1.**
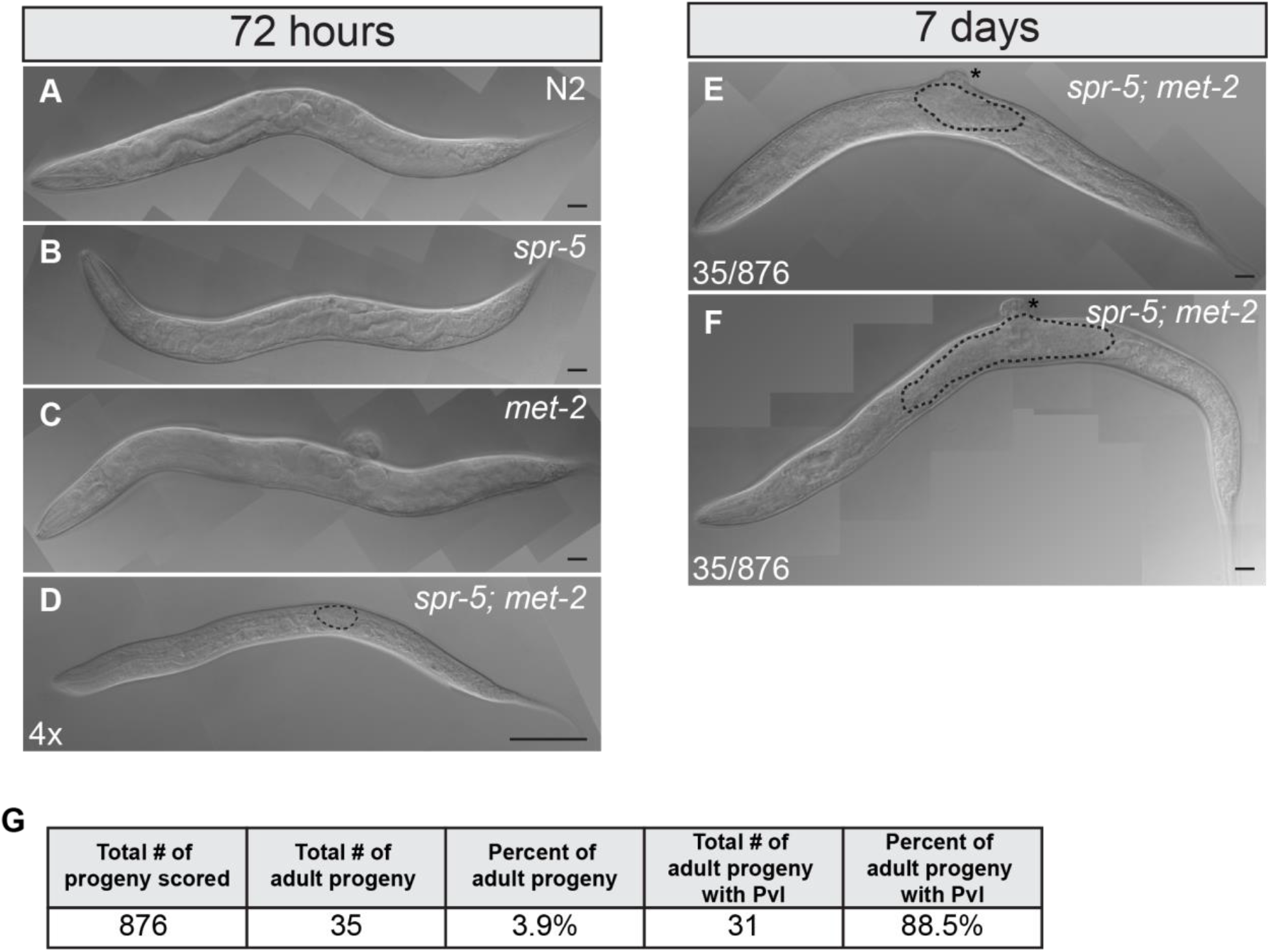
*spr-5; met-2* mutants display severe developmental delay and protruding vulva. 40x DIC images of N2 (A), *spr-5* (B), *met-2* (C), and *spr-5; met-2* (D) progeny at 72 hours post synchronized lay. DIC image in (D) was magnified an additional 4x. 40x DIC images of two examples of *spr-5; met-2* adult progeny (E, F) at seven days post synchronized lay. Dashed-line (D) outlines germline and asterisks (E, F) denote protruding vulva. Scale bar: 100μm. (G) Quantification of *spr-5; met-2* progeny that reached the adult stage by seven days post synchronized lay, along with quantification of protruding vulva (Pvl) in these animals that reached the adult stage. The 876 progeny scored came from a total of 25 hermaphrodites across two independent experiments.

**Figure 2 - supplement figure 1.**
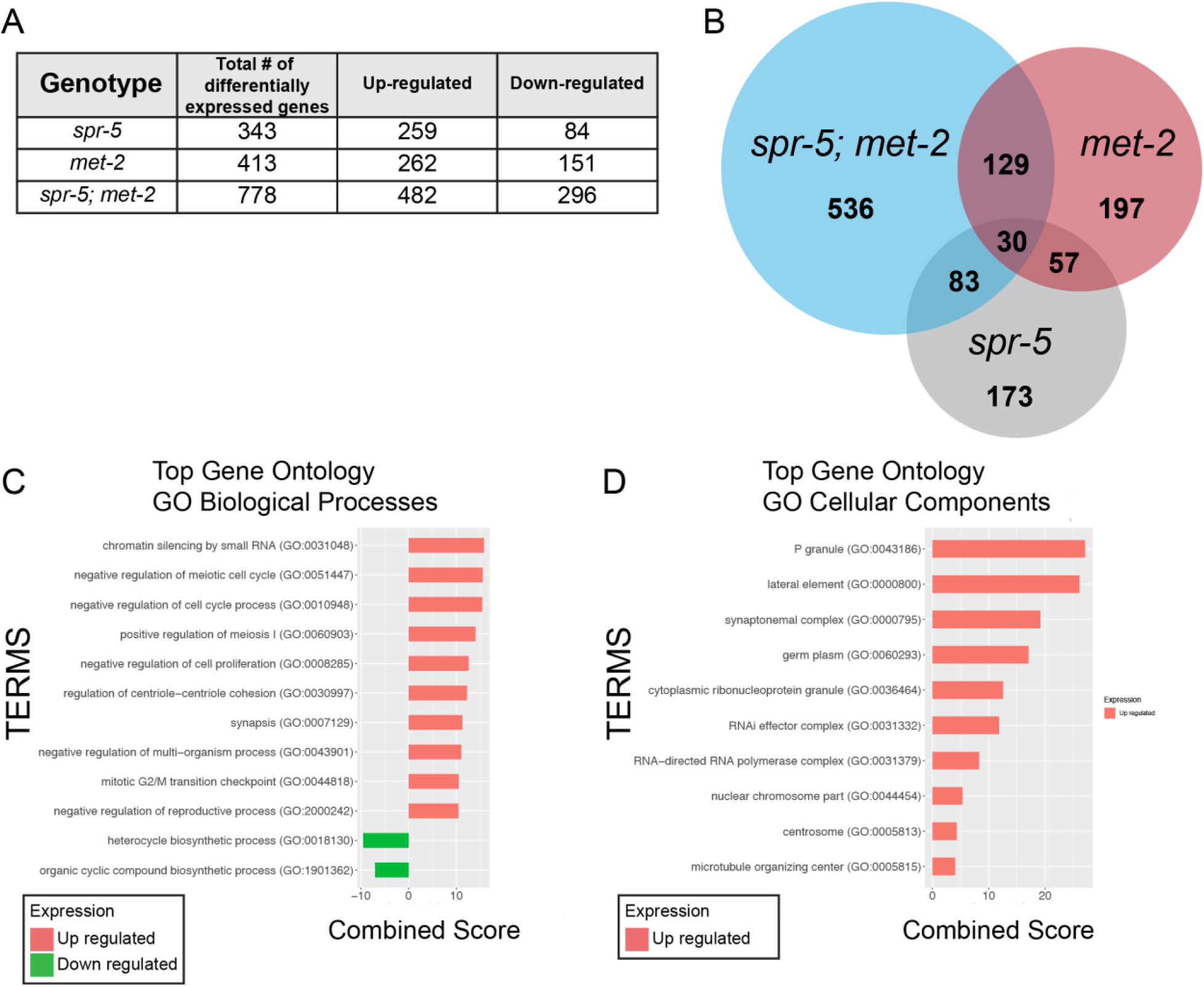
Differential gene expression in *spr-5, met-2, and spr-5; met-2* progeny *compared* to N2 progeny. (A) Table summary of differentially expressed genes in *spr-5, met-2, and spr-5; met-2* L1 progeny from DESEQ2 analysis (significance cut-off of p-value < 0.05). (B) Overlap of differentially expressed genes between *spr-5, met-2, and spr-5; met-2* L1 progeny. Gene Ontology analysis showing Biological Processes (C) and Cellular Components (D) amongst genes that were up-regulated (red) and down-regulated (green) in *spr-5; met-2* L1 progeny compared to N2. Combined Score was computed to determine gene set enrichment (Chen et al., 2013) (see source data file 1 for gene ontology R scripts).

**Figure 2 - supplement figure 2.**
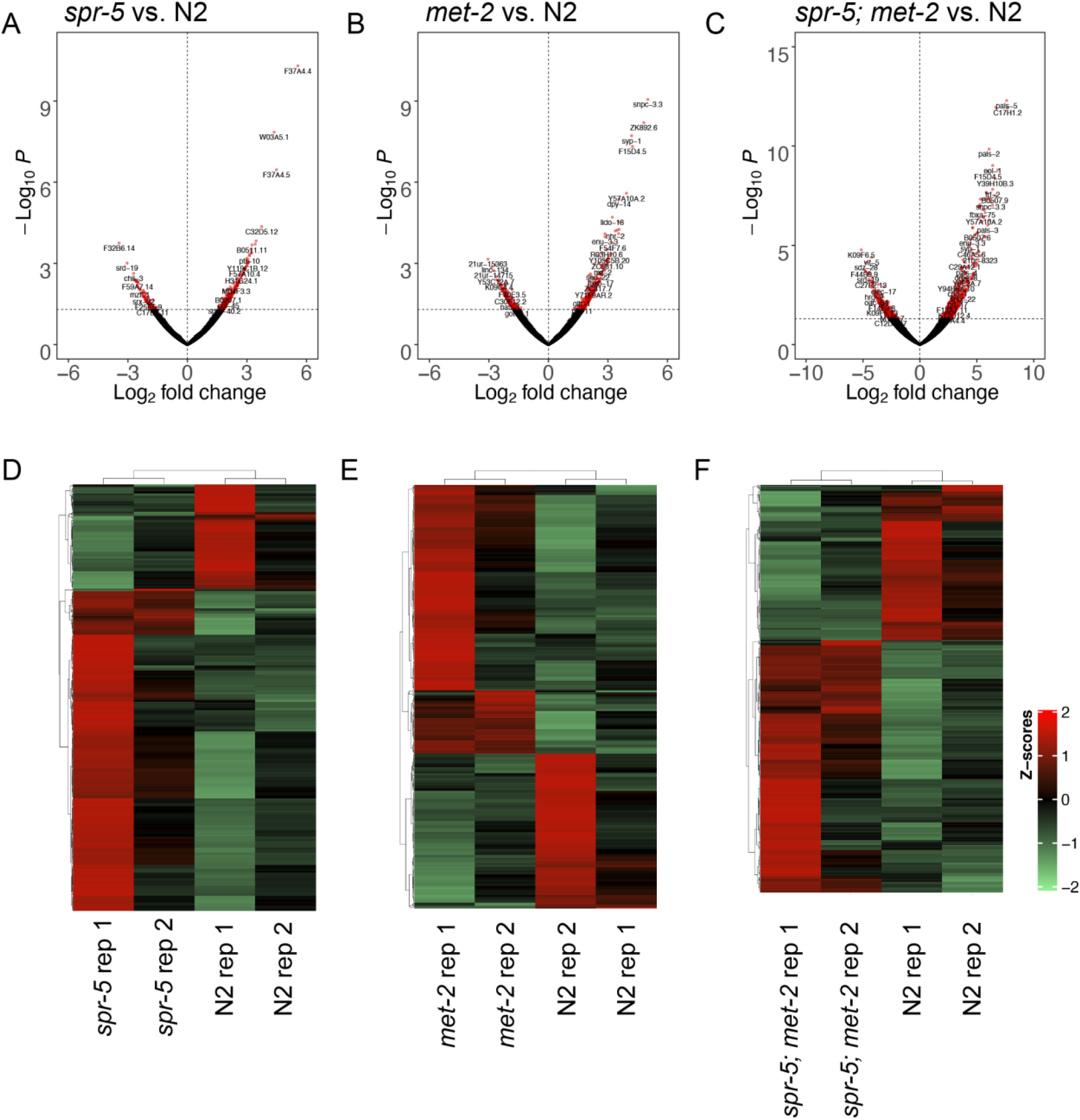
Differential expression and replicate comparison of RNAseq experiments performed on N2, *spr-5*; *met-2*, and *spr-5; met-2* L1 progeny. Volcano plot of log2 fold changes in gene expression (x-axis) by statistical significance (-Log10 P-value; y-axis) in *spr-5* (A), *met-2* (B), and *spr-5; met-2* (C) L1 Progeny compared to N2 (see source data file 2 for volcano plot R scripts). Heatmap of differentially expressed RNA-seq transcripts between N2 and *spr-5* (D), *met-2* (E), and *spr-5; met-2* (F). Data was scaled and hierarchical clustering was performed using the complete linkage algorithm. Distance was measured by calculating pairwise distance. Higher (red) and lower (green) expression is reported as a z-score. (see source data file 3 for heatmap R scripts).

**Figure 2-supplemental figure 3.**
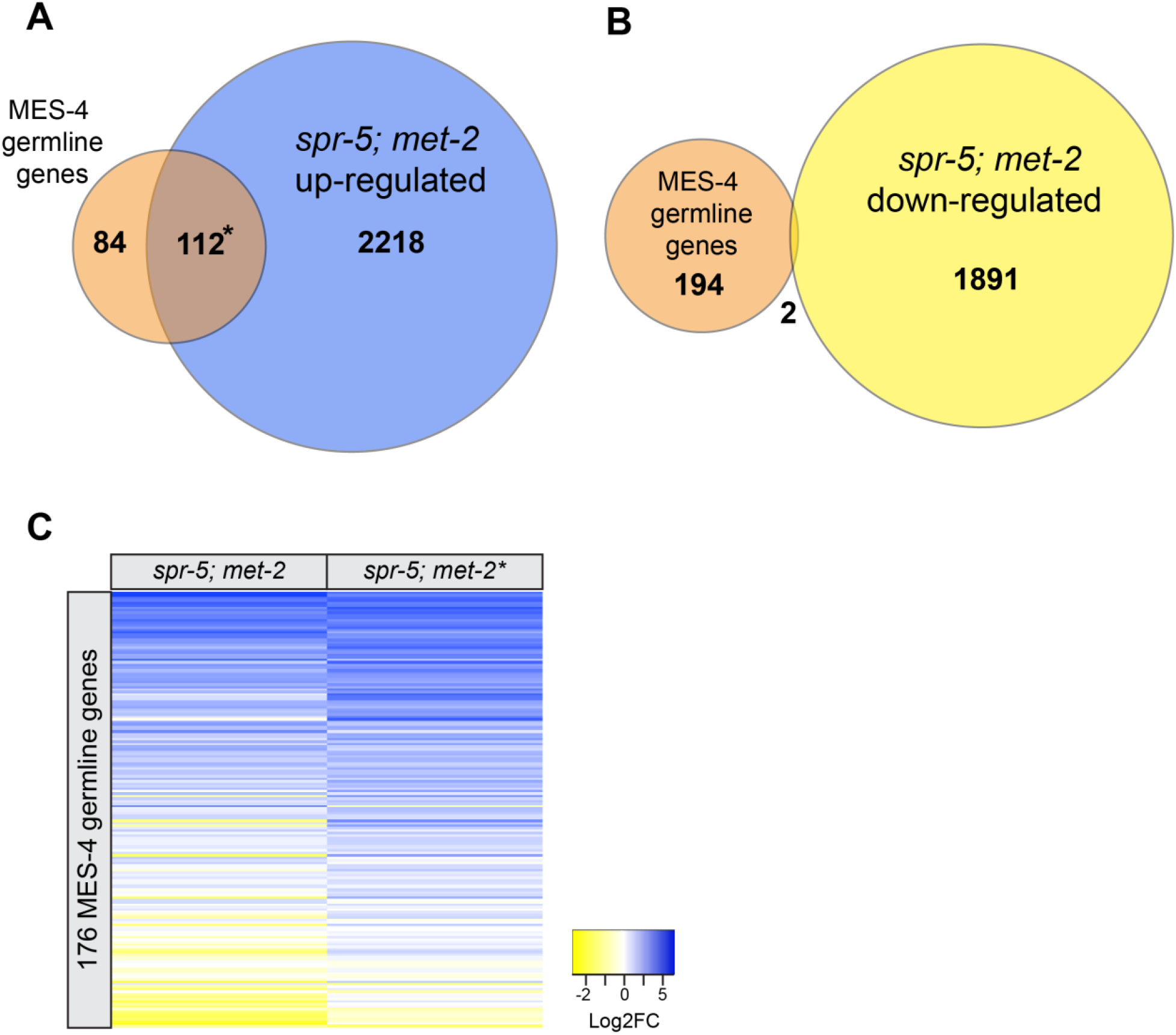
MES-4 germline genes are ectopically expressed in *spr-5; met-2* mutant soma in the RNAseq repeat experiment. Overlap between MES-4 germline genes and genes up-regulated (A) and down-regulated (B) in *spr-5; met-2* L1 progeny. Significant over-enrichment in A was determined by the hypergeometric test (*P-value < 1.20E-54). (C) Heatmap of log2 fold change (FC) of 176 MES-4 germline genes in *spr-5; met-2* mutants compared to N2 (2 replicates, low-input, see methods) vs. repeat of *spr-5; met-2\** mutants compared to N2 (*3 replicates, standard Poly-A selection, see methods). Log2FC values are represented in a yellow to blue gradient and range from −2 to 5 and were sorted by the average log2FC in *spr-5; met-2* and *spr-5; met-2\** progeny. Yellow represents genes with negative log2FC values and blue represents genes with positive log2FC values compared to N2. The remaining 21 MES-4 germline genes were not included because they do not have an expression value in one or more of the data sets (*spr-5; met-2* or *spr-5; met-2\**).

**Figure 3 - supplement figure 1.**
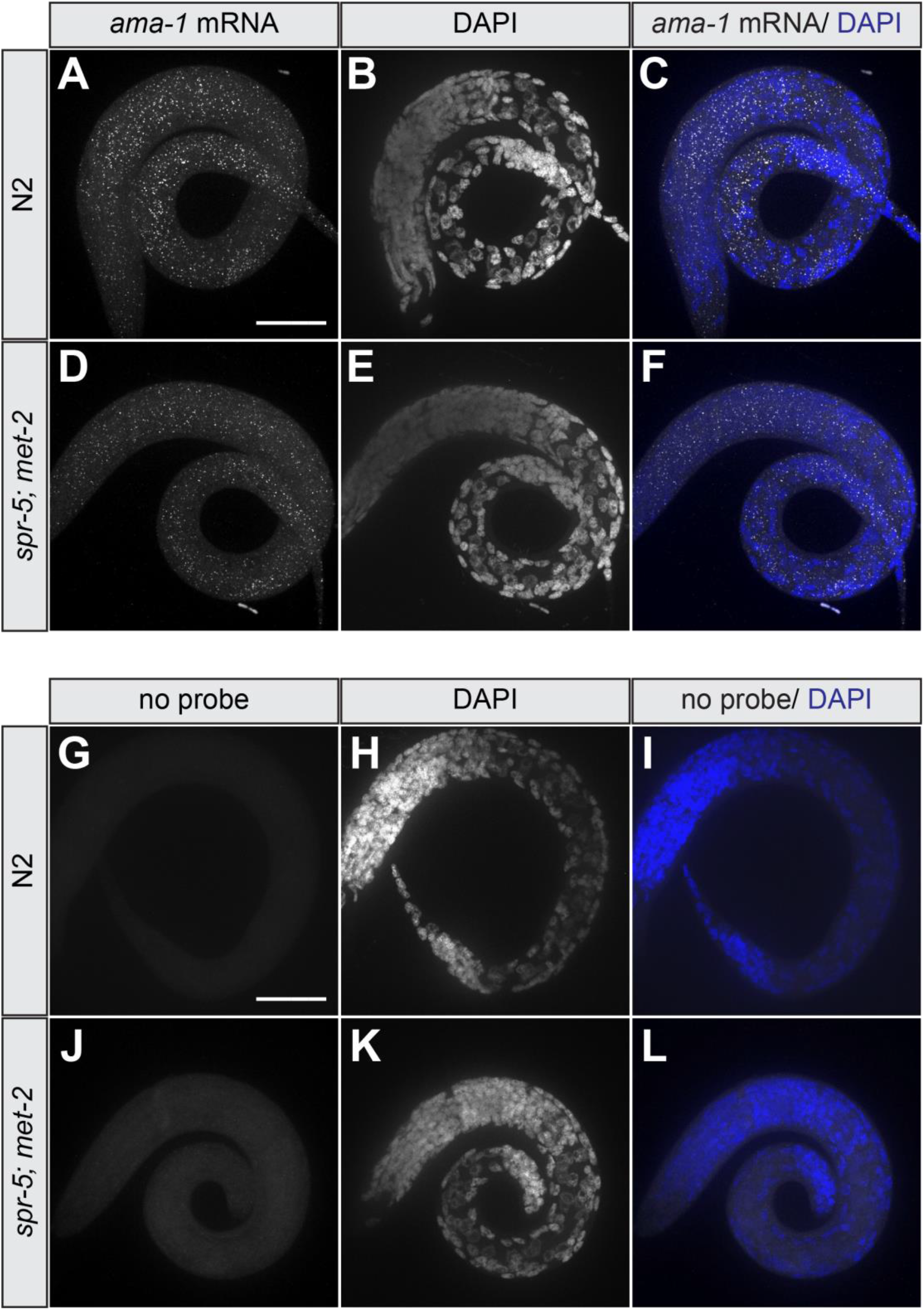
smFISH control experiments. smFISH images of *ama-1* mRNA (A, C, D, F) and a no probe control (G, I, J, L) in N2 (A-C, G-I) and *spr-5; met-2* (D-F, J-L) L1 progeny. DAPI was used as a nuclear marker (B, C, E, F, H, I, K, L). Scale bar 40μm.

**Figure 4 - supplement figure 1.**
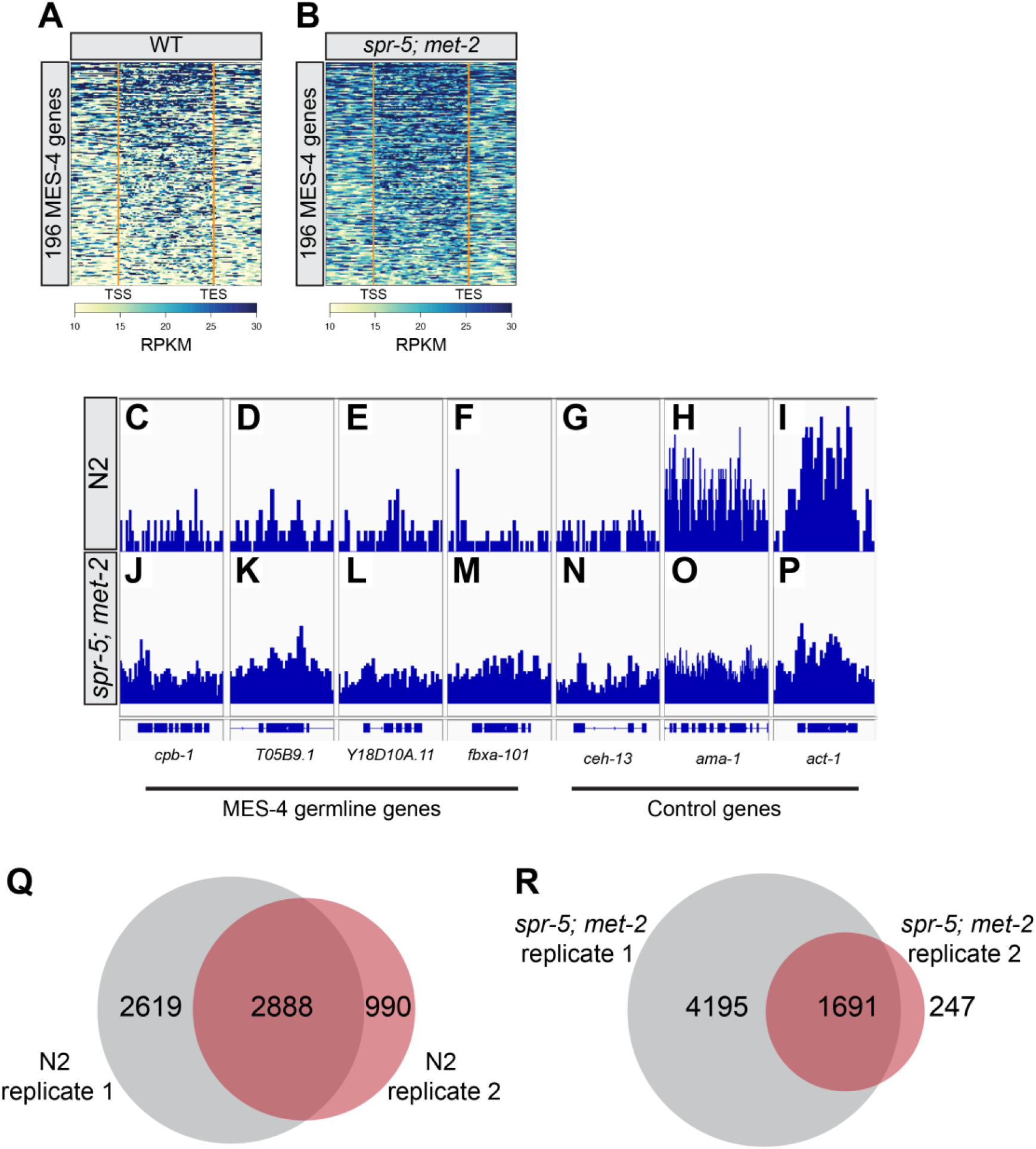
Replicate comparison of H3K36me3 ChIP-seq analysis in *spr-5; met-2* L1 progeny. Heatmap of H3K36me3 ChIP-seq reads from a second replicate (1st replicate in Figure 4) normalized to reads per kilobase million (RPKM) over the gene bodies of 196 MES-4 germline genes in wild-type (N2) (A) versus *spr-5; met-2* (B) L1 progeny. Gene bodies were pseudoscaled to 1kb with 500bp borders separated by orange bars that represent the transcriptional start site (TSS) and transcriptional end site (TES). Integrative Genome Viewer (IGV) image of H3K36me3 ChIP-seq reads from replicate 2 normalized to RPKM at MES-4 germline genes (C-F, J-M) versus control genes (G-I, N-P) in N2 (C-I) and *spr-5; met-2* (J-P) L1 progeny. RPKM IGV windows were scaled between 0 and 107 RPKM for all genes. Venn-diagram displaying the overlap of H3K36me3 ChIP-seq called broad peaks for N2 (Q) and *spr-5; met-2* (R) replicates.

**Figure 4 - supplement figure 2.**
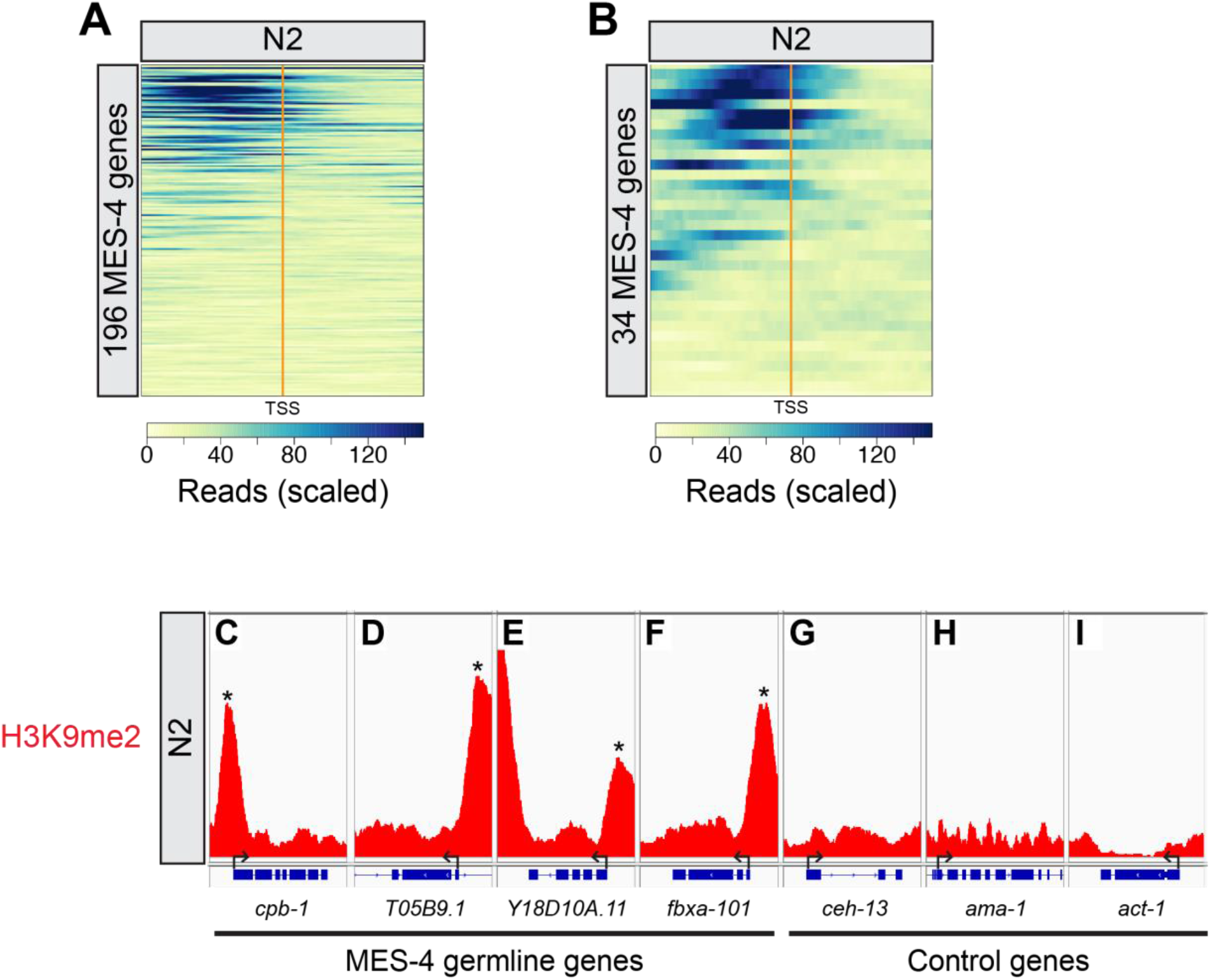
MES-4 germline genes display H3K9me2 at their promoters. Heatmap of H3K9me2 promoter peaks in N2 L1 progeny at all 196 MES-4 germline genes (A), and at the 34 MES-4 germline genes that were ectopically expressed in the soma of *spr-5; met-2* progeny (B). ChIP-seq reads were scaled to genome wide coverage for H3K9me2 (15 million reads). The transcriptional start site (TSS) is denoted by an orange bar with 500bp flanking regions upstream and downstream of the TSS. Integrative Genome Viewer (IGV) image of ChIP-seq reads from N2 L1 progeny scaled to genome wide coverage for H3K9me2 (15 million reads) over the promoters of MES-4 germline genes that were ectopically expressed in the soma of *spr-5; met-2* progeny (C-F) and control genes (G-I). IGV windows were scaled between 0 and 255 RPKM. Asterisks (*) denotes H3K9me2 promoter peaks at MES-4 germline genes.

**Supplementary file 1. Raw scores for developmental delay in N2, *spr-5*, *met-2*, and *spr-5; met-2* progeny.** Progeny were scored 72 hours after a synchronized lay as developing to the L1, L2, L3, L4, or young adult (YA) stage (Figure 1A-E). N= the total number of progeny from a total of 20-25 hermaphrodites scored over three experiments.

**Supplementary file 2. Differentially expressed transcripts in *spr-5*, *met-2*, and *spr-5; met-2* progeny.** List of differentially expressed transcripts in *spr-5*, *met-2*, *spr-5; met-2* (Wald test, P-value<0.05), and the repeat RNA-seq experiment with three additional *spr-5; met-2* replicates (Wald test, P-adj<0.05) L1 progeny compared to N2 progeny from DESEQ2 analysis (Figure 2-supplement figure 2A-D).

**Supplementary file 3. 176 MES-4 germline gene log2(FC) values used to generate heatmap.** The log2(FC) values of the 176 MES-4 germline genes from the DESEQ2 analysis performed on *spr-5, met-2, spr-5; met-2*, and the repeat RNA-seq experiment of *spr-5; met-2* (*) L1 progeny compared to N2 progeny. log2(FC) values were sorted from highest to lowest based on the differential expression of genes in *spr-5; met-2* progeny compared to N2 (Figure 2C-D).

**Supplementary file 4. smFISH probe sequences.** List of smFISH probe sequences for *htp-1* and *cpb-1* used in smFISH experiments (Figure 3).

**Supplementary file 5. Primer sequences used for genotyping and RT-PCR.** List of genotyping primers and RT-PCR primers (Figure 5).

**Supplementary file 6. Raw values from Quantitative RT-PCR analysis.** Raw SQ means and standard deviations for RT-PCR experiments (Figure 5).

**Source data file 1**. R scripts for gene ontology (Figure 2-supplement figure 1C-D)

**Source data file 2**. R scripts for volcano plots (Figure 2-supplement figure 2A-C)

**Source data file 3**. R scripts for heatmaps (Figure 2-supplement figure 2D-F)

