## Supplemental Data 7 for "*C. elegans* establishes germline versus soma by balancing inherited histone methylation"

```
library(tidyverse)
library(ggplot2)

setwd("/users/admin/dropbox/datasets/Brandon/GO")

#read in GO Biologcial Process from WormEnrichr

setwd("/users/admin/dropbox/datasets/Brandon/GO/SPR5MET2.UP")

enriched.up  <- read.delim("GO_Biological_Process_2018_table.txt", header = TRUE, sep = "\t", dec = ".")

setwd("/users/admin/dropbox/datasets/Brandon/GO/SPR5MET2.DOWN")

enriched.down <- read.delim("GO_Biological_Process_2018_table.txt", header = TRUE, sep = "\t", dec = ".")

up <- filter(enriched.up, P.value < 0.05)
down <- filter(enriched.down, P.value < 0.05)

up$type <- "up"
down$type <- "down"
up <- up [c(1:10),]
up <- up[order(up$Combined.Score), ]  # sort
down <- down [c(1:2),] ## only 2 ontologies significant
down <- down[order(down$Combined.Score), ]  # sort
down$Combined.Score <- (-1) * down$Combined.Score
gos <- rbind(down,up)
gos$Term <- factor(gos$Term, levels=gos$Term)
# Diverging Barcharts
g <- ggplot(gos, aes(x=Term, y=Combined.Score , label=Combined.Score)) + 
  geom_bar(stat='identity', aes(fill=type), width=.5,position="dodge")  +
  scale_fill_manual(name="Expression", 
                    labels = c("Down regulated", "Up regulated"), 
                    values = c("down"="#00ba38", "up"="#f8766d")) + 
  labs(subtitle="Combined scores (GO Biological Process)", 
       title= "Top Gene Ontologies") + 
  coord_flip()
g+ theme(axis.text=element_text(size=17),
          axis.title=element_text(size=15),
          plot.title = element_text(size=22),
          plot.subtitle = element_text(size = 11))


#read in GO Cellular Component from WormEnrichr

setwd("/users/admin/dropbox/datasets/Brandon/GO/SPR5MET2.UP")

enriched.up <- read.delim("GO_Cellular_Component_2018_table.txt", header = TRUE, sep = "\t", dec = ".")

setwd("/users/admin/dropbox/datasets/Brandon/GO/SPR5MET2.DOWN")

enriched.down<- read.delim("GO_Cellular_Component_2018_table.txt", header = TRUE, sep = "\t", dec = ".")

up <- filter(enriched.up, P.value < 0.05)
down <- filter(enriched.down, P.value < 0.05)

up$type <- "up"
up <- up [c(1:10),]
up <- up[order(up$Combined.Score), ]  # sort
gos <- rbind(down,up)
gos$Term <- factor(gos$Term, levels=gos$Term)
# Diverging Barcharts
g <- ggplot(gos, aes(x=Term, y=Combined.Score , label=Combined.Score)) + 
  geom_bar(stat='identity', aes(fill=type), width=.5,position="dodge")  +
  scale_fill_manual(name="Expression", 
                    labels = c("Up regulated"), 
                    values = c( "up"="#f8766d")) + 
  labs(subtitle="Combined scores (GO Cellular Component)", 
       title= "Top Gene Ontologies") + 
  coord_flip()
g+ theme(axis.text=element_text(size=17),
         axis.title=element_text(size=15),
         plot.title = element_text(size=22),
         plot.subtitle = element_text(size = 11))
```
