## Supplemental Data 8 for "*C. elegans* establishes germline versus soma by balancing inherited histone methylation"

```
library(ggrepel)
library(EnhancedVolcano)

setwd("/Users/admin/Dropbox/Datasets/Brandon/")

#read in deseq2 output files (NA values removed)
SPR5.na <- read.delim("SPR5.rmna.txt", header = TRUE, sep = "\t", dec = ".")
SPR5MET2.na <- read.delim("SPR5MET2.rmna.txt", header = TRUE, sep = "\t", dec = ".")
MET2.na <- read.delim("MET2.rmna.txt", header = TRUE, sep = "\t", dec = ".")

##Draw volcano plots using enhanced volcano package

#Volcano plot for SPR5
plot <- EnhancedVolcano( SPR5.na, 
                          lab = SPR5.na$symbol,
                          x = "log2fc",
                          y = "p.value",
                          pCutoff = 0.05, 
                          FCcutoff = 0,
                          xlab = bquote(~Log[2]~ "fold change"),
                          ylab = bquote(~-Log[10]~italic(P)),
                          axisLabSize = 40,
                          transcriptPointSize = 2,
                          transcriptLabSize = 5,
                          title = "SPR5 vs N2",
                          titleLabSize = 40,
                          col = c("black", "black", "black", "red3"),
                          colAlpha = 0.5,
                          xlim = c(-6,6),
                          ylim = c(0,11),
                          cutoffLineType = "dashed",
                          cutoffLineCol = "black",
                          cutoffLineWidth = 0.5,
                          legendPosition = "none",
                          DrawConnectors = FALSE,
                          widthConnectors = 0.2,
                          colConnectors = "grey30",
                          border = "full",
                          borderWidth = 1.5,
                          borderColour = "black",
                          gridlines.major = FALSE,
                          gridlines.minor = FALSE)

#Volcano plot for MET2
plot2 <- EnhancedVolcano( MET2.na, 
                         lab = MET2.na$symbol,
                         x = "log2fc",
                         y = "p.value",
                         pCutoff = 0.05, 
                         FCcutoff = 0,
                         xlab = bquote(~Log[2]~ "fold change"),
                         ylab = bquote(~-Log[10]~italic(P)),
                         axisLabSize = 40,
                         transcriptPointSize = 2,
                         transcriptLabSize = 5,
                         title = "MET2 vs N2", 
                         titleLabSize = 40,
                         col = c("black", "black", "black", "red3"),
                         colAlpha = 0.5,
                         xlim = c(-6,6),
                         ylim = c(0,11),
                         cutoffLineType = "dashed",
                         cutoffLineCol = "black",
                         cutoffLineWidth = 0.5,
                         legendPosition = "none",
                         DrawConnectors = FALSE,
                         widthConnectors = 0.2,
                         colConnectors = "grey30",
                         border = "full",
                         borderWidth = 1.5,
                         borderColour = "black",
                         gridlines.major = FALSE,
                         gridlines.minor = FALSE)

#Volcano plot for SPR5MET2
plot3 <- EnhancedVolcano( SPR5MET2.na, 
                         lab = SPR5MET2.na$symbol,
                         x = "log2fc",
                         y = "p.value",
                         pCutoff = 0.05, 
                         FCcutoff = 0,
                         xlab = bquote(~Log[2]~ "fold change"),
                         ylab = bquote(~-Log[10]~italic(P)),
                         axisLabSize = 40,
                         transcriptPointSize = 2,
                         transcriptLabSize = 5,
                         title = "SPR5MET2 vs N2", 
                         titleLabSize = 40,
                         col = c("black", "black", "black", "red3"),
                         colAlpha = 0.5,
                         xlim = c(-10,10),
                         ylim = c(0,15),
                         cutoffLineType = "dashed",
                         cutoffLineCol = "black",
                         cutoffLineWidth = 0.5,
                         legendPosition = "none",
                         DrawConnectors = FALSE,
                         widthConnectors = 0.2,
                         colConnectors = "grey30",
                         border = "full",
                         borderWidth = 1.5,
                         borderColour = "black",
                         gridlines.major = FALSE,
                         gridlines.minor = FALSE)


                 
```
