## Supplemental Data 9 for "*C. elegans* establishes germline versus soma by balancing inherited histone methylation"

```
library(RColorBrewer)
library(tidyverse)
library(ComplexHeatmap)
library(circlize)

#set working directory to where deseq2 output files are located
setwd("/Users/admin/Dropbox/Datasets/Brandon/deseq")

#read in deseq2 output files
MET2.counts <- tbl_df(read.delim("Galaxy204-[Normalized_counts_file_on_MET2_vs_N2].tabular",
                                 header = TRUE, sep = "\t", stringsAsFactors = FALSE))

SPR5.counts <- tbl_df(read.delim("Galaxy195-[Normalized_counts_file_on_SPR5_vs_N2].tabular",
                                 header = TRUE, sep = "\t", stringsAsFactors = FALSE))

SPR5MET2.counts <- tbl_df(read.delim("Galaxy186-[Normalized_counts_file_on_SPR5MET2_vs_N2_DESEQ2].tabular",
                                     header = TRUE, sep = "\t", stringsAsFactors = FALSE))

#change column header names
colnames(MET2.counts)  <- c("geneid", "N22", "N21","MET22", "MET21" )
colnames(SPR5.counts)  <- c("geneid", "N22", "N21","SPR52", "SPR51" )
colnames(SPR5MET2.counts)  <- c("geneid", "N22", "N21","SPR5MET21", "SPR5MET22" )

#change working directory  
setwd("/Users/admin/Dropbox/Datasets/Brandon/")

#read in csv files containing differentially expressed genes
MET2.DEGs <- tbl_df(read.csv("MET2.DEGs.csv", stringsAsFactors = FALSE))
SPR5.DEGs <- tbl_df(read.csv("SPR5.DEGs.csv", stringsAsFactors = FALSE))
SPR5MET2.DEGs <- tbl_df(read.csv("SPR5MET2.DEGs.csv", stringsAsFactors = FALSE))

#subset normalized counts files to differentially expressed genes
met2.heat <- MET2.counts %>% 
  inner_join(MET2.DEGs, by = "geneid") %>% 
  arrange(p.value) 

spr5.heat <- SPR5.counts %>% 
  inner_join(SPR5.DEGs, by = "geneid") %>% 
  arrange(p.value) 

spr5met2.heat <- SPR5MET2.counts %>% 
  inner_join(SPR5MET2.DEGs, by = "geneid") %>% 
  arrange(p.value) 

# set colors for heat maps
col_fun = colorRamp2(c(-2, 0, 2), c("light green", "black", "red"))

# Heatmaps
# log2(normalized counts) of differntially expressed genes
# ComplexHeatmap package

## Met2 heatmap
met2 <- met2.heat %>% 
  mutate("N2 rep 2"= log2(N22 + 1), "N2 rep 1"= log2(N21 +1), #log2 transformation
         "met-2 rep 2"= log2(MET22 +1), "met-2 rep 1"= log2(MET21 +1)) %>% 
  select("N2 rep 1", "N2 rep 2", "met-2 rep 1", "met-2 rep 2") %>% #filter out everything except log transformed counts
  data.matrix() # transform dataframe to matrix
  
met2_scaled = t(scale(t(met2))) #perform scaling
#create heatmap with complexheatmap package
Heatmap(met2_scaled,
        name = "met-2 scaled",
        col = col_fun,
        cluster_rows = TRUE,
        cluster_columns = TRUE,
        clustering_method_rows = "complete",
        clustering_method_columns = "complete",
        clustering_distance_rows = function(x, y) 1 - cor(x, y),  #function to calculate pairwise distance
        clustering_distance_columns = function(x, y) 1 - cor(x, y),
        heatmap_legend_param = list (
          title = 'Z-scores',
          legend_height = unit(4, "cm"),
          title_position = "lefttop-rot"
        ))

## spr5 heatmap

spr5 <- spr5.heat %>% 
  mutate("N2 rep 2"= log2(N22 + 1), "N2 rep 1"= log2(N21 +1), 
         "spr-5 rep 2"= log2(SPR52 +1), "spr-5 rep 1"= log2(SPR51 +1)) %>% 
  select("N2 rep 1", "N2 rep 2", "spr-5 rep 1", "spr-5 rep 2") %>% 
  data.matrix()

spr5_scaled = t(scale(t(spr5)))
Heatmap(spr5_scaled,
        name = "spr5 scaled",
        col = col_fun,
        #column_order = sort(colnames(spr5_scaled)),
        cluster_rows = TRUE,
        cluster_columns = TRUE,
        clustering_method_rows = "complete",
        clustering_method_columns = "complete",
        clustering_distance_rows = function(x, y) 1 - cor(x, y),
        clustering_distance_columns = function(x, y) 1 - cor(x, y),
        heatmap_legend_param = list (
          title = 'Z-scores',
          legend_height = unit(4, "cm"),
          title_position = "lefttop-rot"
        ))

## spr5-met2 heatmap

spr5met2 <- spr5met2.heat %>% 
  mutate("N2 rep 2"= log2(N22 + 1), "N2 rep 1"= log2(N21 +1), 
         "spr-5;met-2 rep 1"= log2(SPR5MET21 +1), "spr-5;met-2 rep 2"= log2(SPR5MET22 +1)) %>% 
  select("N2 rep 1", "N2 rep 2", "spr-5;met-2 rep 1", "spr-5;met-2 rep 2") %>% 
  data.matrix()

spr5met2_scaled = t(scale(t(spr5met2)))

Heatmap(spr5met2_scaled,
        name = "spr5met2 scaled",
        col = col_fun,
        #column_order = sort(colnames(spr5_scaled)),
        cluster_rows = TRUE,
        cluster_columns = TRUE,
        clustering_method_rows = "complete",
        clustering_method_columns = "complete",
        clustering_distance_rows = function(x, y) 1 - cor(x, y),
        clustering_distance_columns = function(x, y) 1 - cor(x, y),
        heatmap_legend_param = list (
          title = 'Z-scores',
          legend_height = unit(4, "cm"),
          title_position = "lefttop-rot"
        ))
```
